# Thermodynamic, Kinetic, and Structural Determinants of Ligand Selectivity in A2A and A2B Adenosine Receptors

**DOI:** 10.64898/2026.08.24.746716

**Authors:** Omar Alsina, Samuele Di Cristofano, Stefano Raniolo, Vittorio Limongelli

## Abstract

G protein-coupled receptors are major pharmacological targets, yet achieving subtype selectivity remains challenging when closely related receptors share highly conserved orthosteric binding sites. Here, we investigate the molecular determinants governing ligand recognition and unbinding at the adenosine A2A and A2B receptors, two closely related class A GPCRs with markedly different pharmacological profiles. We combine Funnel Metadynamics and adaptive infrequent metadynamics to characterize the thermodynamics and kinetics of three representative ligands: the non-selective antagonist theophylline, the A2A-selective inverse agonist ZM-241385, and the non-selective full-agonist NECA. Across six ligand–receptor complexes, our simulations reproduce experimentally resolved binding modes, predict the unresolved binding poses of TEP and ZMA at A2B, and provide binding free energies consistent with experimental trends. Kinetic simulations further resolve ligand-specific unbinding pathways, metastable intermediates, residence times, rate-determining transitions, and their associated transition-state configurations. Comparison of A2A and A2B reveals how subtle differences within and around their highly conserved orthosteric sites are amplified into distinct thermodynamic and kinetic behaviors. In particular, we identify three major receptor-specific features: differences in hydration and polarity near TM1/TM2/TM7, differences in steric packing and pocket volume at the TM3/TM5/TM6 floor, and a more dynamic network of charged extracellular residues and lipids in A2B that modulates ligand egress. These features rationalize ligand-dependent differences in affinity, residence time, and subtype selectivity, including the preferential stabilization of ZMA-like antagonists at A2A. Overall, our results provide a dynamic atomistic map of the A2A and A2B orthosteric regions and demonstrate how thermodynamic and kinetic information can reveal pharmacologically relevant differences that are not apparent from static structures alone. This framework may support the rational design and repurposing of subtype-selective adenosine receptor ligands.

## Introduction

G protein-coupled receptors (GPCRs) constitute one of the largest families of membrane signalling proteins and regulate a broad spectrum of physiological processes by translating extracellular chemical signals into intracellular responses.^1–3^ Their activation modulates diverse signalling pathways, including cyclic adenosine monophosphate (cAMP) production, intracellular calcium mobilisation, gene expression, and ion transport.^4^ Accordingly, dysregulation of GPCR signalling is associated with numerous human diseases with significant social and economic consequences, including metabolic, neurological, cardiovascular, and oncological disorders.^5^ For this reason, GPCRs have gained the attention of pharmacological research in recent years, and as of today, approximately 35% of all FDA-approved drugs target a GPCR.^6–8^ In this paper, we focus on the adenosine receptors (ARs) 2A (A2A) and 2B (A2B), which are class A GPCRs that are predominantly expressed in the cardiovascular, nervous, and immune tissues.^9^ They play a central role in diseases such as obesity, cardiovascular diseases, Parkinson’s disease, and cancer.^10–14^ Despite having a sequence identity of 58% across the whole chain and a highly conserved orthosteric binding site (OBS) (84 % sequence identity), except for a handful of point mutations in the transmembrane (TM) helix 6, TM 7, and ECL3, A2B exhibits low affinity for the endogenous agonist adenosine.^15–17^ See Figure 1 for a spatial representation of the protein structures, their sequence identity, and the OBS point mutations. Therefore, due to the sequence similarities between A2A and A2B receptors, it is extremely challenging to target the desired AR subtype without suffering from off-target binding and potentially inducing undesirable side effects.^18,19^ For example, targeting A2A activation with an agonist can induce tissue protection of peripheral organs, whilst activation of A2B can induce tumor growth in the bladder and breasts.^20–22^ Therefore, if the same agonist activates both receptor subtypes, then a desired A2A activation can lead to catastrophic downstream side effects by off-target A2B activation, or vice versa. Thankfully, to date, there are over 90 Protein Data Bank (PDB) entries for A2A in different activation states with a variety of different ligands bound to its OBS, which elucidate the atomistic features that differentiate the various ligand binding modes and their interactions with the receptor.^23^ A2B, however, does not share the same fortune, with only four entries in the PDB to date, and all of them are in their fully activated state with agonists bound in their OBS; therefore, the experimental structural information we have of A2B is limited to its active state.^24,25^ Despite the large number of elucidated A2A structures, these static representations do not necessarily explain the differences in ligand affinities and kinetics between the receptor subtypes. To rationalise these differences, it is necessary to explore the dynamics of the system at an atomistic resolution.^26–29^ *In vitro* assays cannot elucidate these dynamical processes, but fortunately, molecular dynamics (MD) algorithms are capable of modelling and simulating complex biological processes, such as GPCRs, with a high level of accuracy to approximate ligand-binding processes.^30,31^ In this paper, we make use of an MD-based protocol that is capable of accurately characterizing the thermodynamics and the kinetics of three ligands binding to both A2A and A2B receptors: theophylline (TEP), a non-selective antagonist for both receptors; ZM-241385 (ZMA), an A2A-selective inverse agonist; and NECA, a non-selective full agonist.^32–34^ Our protocol leverages the use of enhanced sampling MD techniques such as Funnel Metadynamics (FM) (developed in our group) and adaptive infrequent metadynamics (AdInf), which, when blended with our expertise on GPCR systems, can delineate minute differences and similarities between the ligands’ binding modes and unbinding pathways that ultimately give rise to thermodynamic and kinetic observations.^35–41^ Importantly, these two enhanced-sampling approaches are inherently complementary: FM provides a detailed equilibrium thermodynamic description of the binding landscape, while AdInf resolves the non-equilibrium dynamical processes governing ligand unbinding, together yielding a unified and internally consistent picture of receptor–ligand recognition. With FM, we compute a free energy surface (FES) for each ligand-protein pair, which not only gives us an estimate of the Boltzmann distribution for all the bound and unbound states but also allows us to compute their binding affinities (△G), which can be directly compared to *in vitro* experiments. Moreover, we accurately reproduce the binding modes reported for TEP, ZMA, and NECA bound to A2A, and for NECA bound to A2B, and also offer a prediction of the binding mode for TEP and ZMA bound to A2B, which, to date, has not been experimentally elucidated. Both the reproduced and predicted binding modes are consistent with their respective experimental counterpart, thereby reinforcing the plausibility of the binding modes and the reliability of the FES estimates. Furthermore, with AdInf, we simulate the unbinding of these ligands from their target protein and compute their residence time (RT) at room temperature, as is done similarly by us but not on GPCRs.^42–45^ Using statistically relevant unbinding pathways, we compute a Markov state-to-state transition matrix *M*, which contains all the state-to-state RT with their associated probabilities, along with the identification of the rate-determining step that governs the kinetics of the system. With this information, we elucidate the ephemeral but crucial high-energy transition state (TS) associated with the system’s rate-determining step for unbinding, which is arguably more correlated than the thermodynamic information with *in vivo* efficacy.^46^ The atomistic elucidation of these ephemeral states no doubt aids the community towards the path of rational drug design and repurposing, to increase target efficacy and optimise the kinetics, with the desired biological effect. In conclusion, in this study we present: (1) the reproduction or prediction of the most stable ligand binding mode, (2) a reliable estimate of ligand binding affinity, (3) an estimate of ligand residence time (RT) together with the identification of key metastable states along the unbinding pathway, and (4) the elucidation of the rate-determining step of the unbinding process. Taken together, these four elements provide a detailed atomistic picture of the determinants underlying the similarities and differences in ligand recognition and unbinding at the closely related A2A and A2B receptors. Our findings lay out a receptor-specific OBS map that highlights physicochemical features that are not seen in static structures or in *in vitro* studies. Our results contribute to a deeper understanding of the complex dynamics governing ligand binding and unbinding at A2A and A2B and provide a framework that may motivate analogous mechanistic studies of other pharmacologically relevant systems.

**Figure 1:**
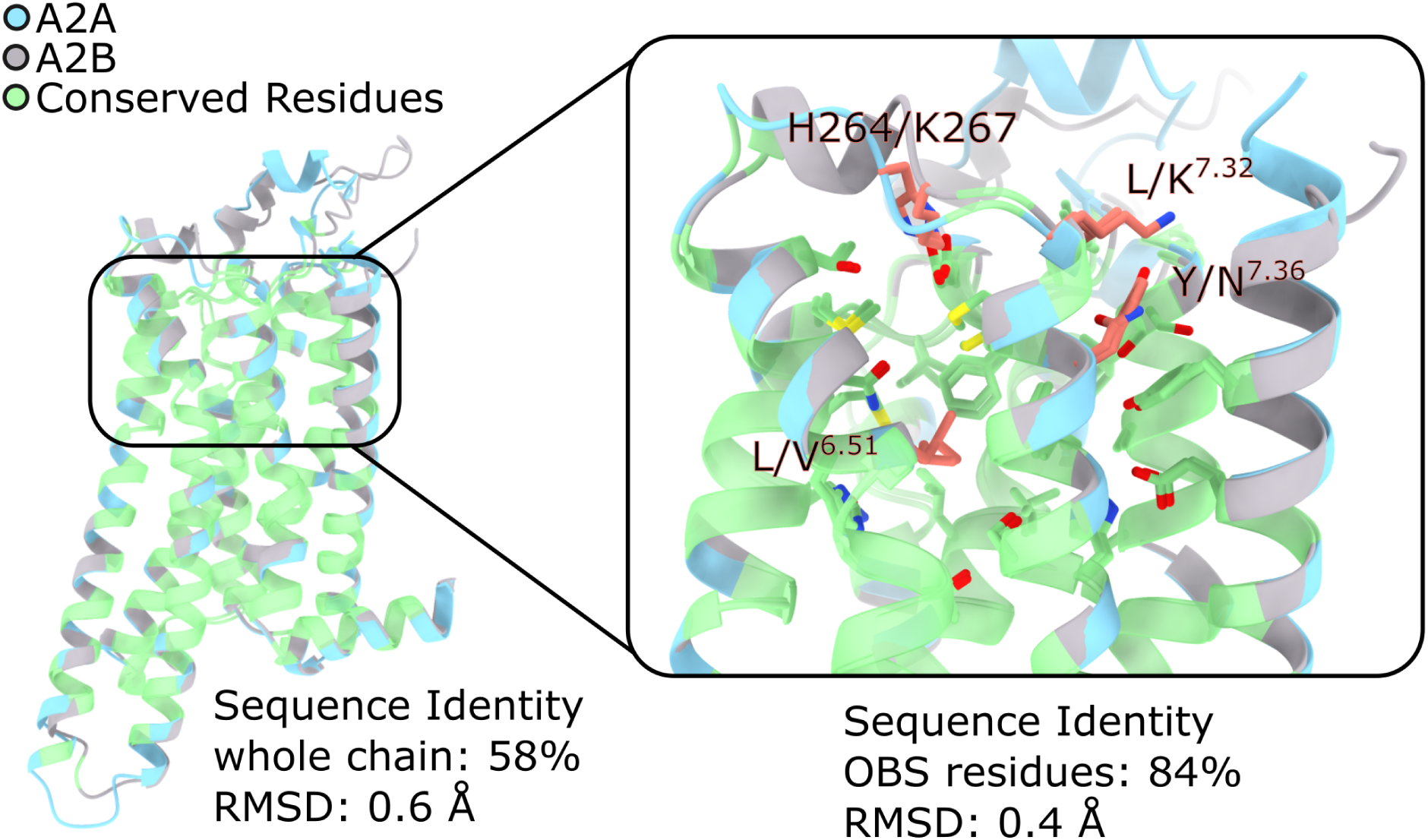
Aligned structural models of A2A (light blue) and A2B (light gray) represented in new cartoon. In green, we show all the residues that are identical between the two receptors. On the left side of the image, we illustrate the high degree of identity that these two receptors show across the whole chain. They exhibit a staggering 58% of sequence identity across residues 1 to 311 and, importantly, these residues sit in an identical spatial arrangement as illustrated by the RMSD of 0.6 Å between the green residues’ *C_α_*. On the right side of the image, we zoom in on the OBS site to illustrate the sequence identity of the main extracellular binding site. The residues forming this pocket are shown in sticks. The sequence identity and the RMSD of *C_α_* atoms are calculated using the shown residues (26 in total), of which only four are different: L/V^6.51^, H264/K267, L/K^7.32^, and Y/N^7.36^, colored in salmon (A2A/A2B, respectively). The identity matrix and RMSD calculation were done with ChimeraX’s Clustal Omega Alignment.

**Figure 2:**
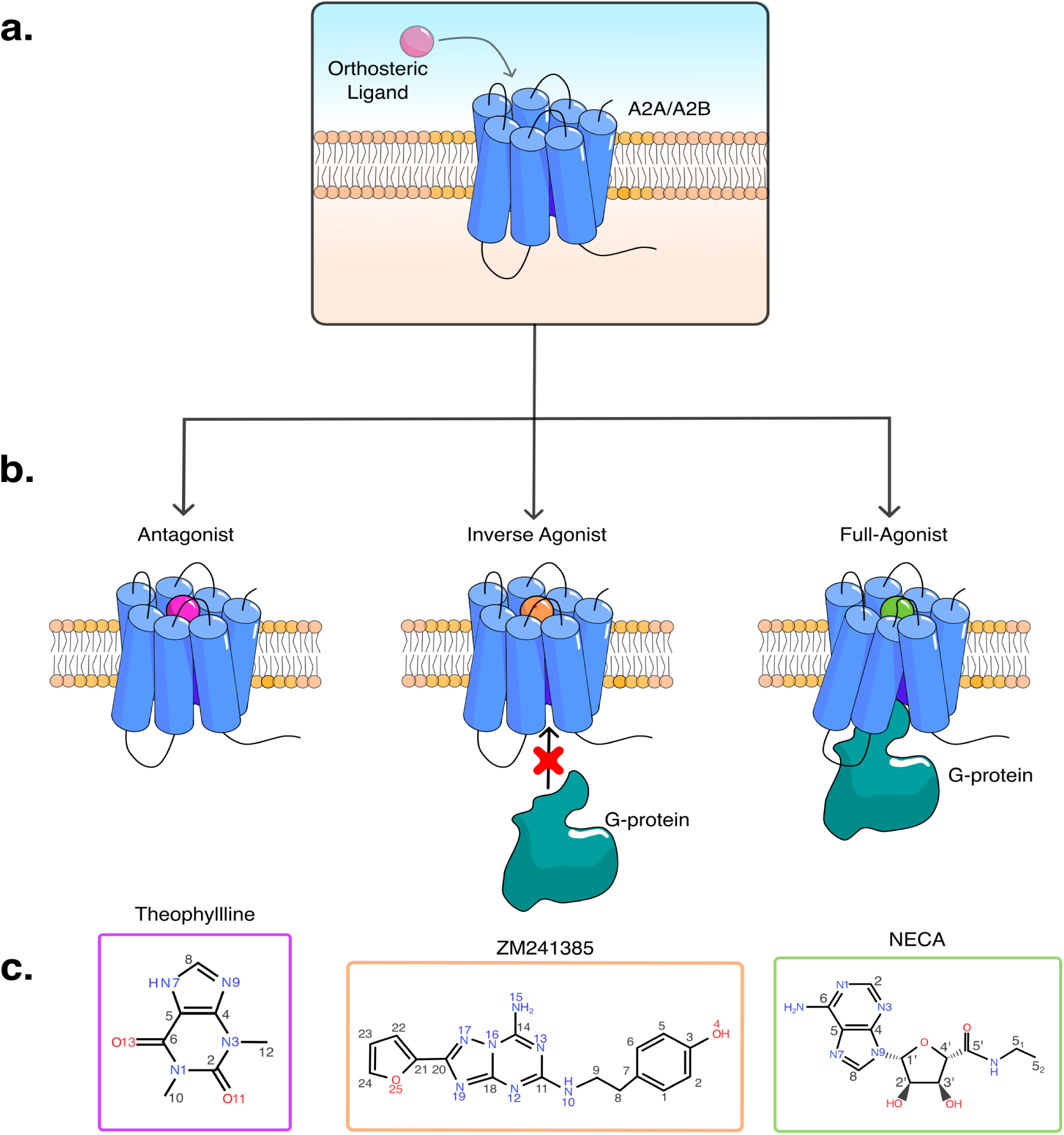
**a** Schematic representation of an A2A/A2B class A GPCR embedded in the cellular membrane. The pink sphere represents a generic ligand bound to the orthosteric binding site. **b** Schematic representation of the three ligands investigated in this study. From left to right, the magenta sphere represents the non-selective antagonist theophylline (TEP); the orange sphere represents the A2A-selective inverse agonist ZM-241385 (ZMA), whose reduction of basal receptor activity is schematically depicted by the absence of G-protein coupling at the intracellular side of the receptor; and the green sphere represents the non-selective full agonist NECA, whose increase in receptor activity is depicted by G-protein coupling. **c** 2D molecular structures of the three ligands, shown using the color coding and heavy-atom numbering adopted throughout this manuscript.

## Methods

### System Setup and unbiased MD simulations

We prepared a total of six ligand-protein complex systems, all with a similar procedure. The A2A structures co-crystallized with the ligand (in parentheses) were taken from the following PDB entries: 5MJZ (TEP), 5IU4 (ZMA), and 5G53 (NECA).^47–49^ The thermostabilizing StaR2-bRIL complex of 5MJZ and 5IU4 was removed, and all the missing parts were reconstructed using MODELLER implemented in the HHpred server.^50–52^ All the mutations present in the models were reverted to their natural wild-type (WT) sequence using UCSF Chimera.^53^ For A2B, the full-active cryo-EM structure of A2B with NECA bound (PDB ID: 7XY7) was used to generate the A2B-NECA-miniG complex, with the miniG protein taken from PDB ID: 5G53. To date, the structure of A2B has only been resolved in its active form through cryo-EM, where the atomic coordinates of extracellular loop 2 (ECL2) are missing.^24^ Hence, we modeled the inactive state of A2B for the TEP-A2B and ZMA-A2B complexes, and we rebuilt the ECL2. The inactive *apo* form deposited in the AlphaFold Protein Structure Database (https://alphafold.ebi.ac.uk/entry/AF-P29275-F1) was used as the starting structure for unbiased MD and replica exchange with solute tempering (REST2) simulations to extensively sample the protein conformational space and refine the ECL2 loop (see “Modelling of the Extracellular Loop 2 in A2B” section in S.I. Text).^54–57^ For inactive A2B complexes, TEP from the PDB ID: 5MJZ was grafted into the OBS after aligning the structures on the C*_α_* atoms of the OBS residues. The same procedure was adopted to manually dock ZMA from PDB ID: 5IU4 to the inactive A2B structure. For both receptors, the N- and C-termini were neutralized by acetylation (ACE) and N-methylamidation (NME), respectively. The protonation states for all residues were selected based on PropKa’s pKa prediction.^58^ All histidines were modeled in the N*ɛ*-tautomer, except for H264 in A2A, which is protonated, and H^7.43^, which is in the N*δ*-tautomeric state. The protonation of H264 is of particular importance for this work. The majority of pKa prediction tools place H264 around 7 - 7.4, hence creating a point of discussion for simulations performed at physiological pH. Previous discussions have suggested that this histidine is neutral at physiological pH values, and the close interaction between H264 and E169 on various crystals is due to the low pH of the crystallization procedure.^59^ However, recent mutagenesis and NMR studies have suggested that this H264 is protonated and forms a salt bridge with E169, having an effect on the ligand’s affinity and kinetics.^60,61^ We have to define H264 in a protonated state, although we recognize the dichotomy that exists around this residue, and we will consider future work on the effects of a neutral H264 on ligand binding in A2A. The protein was then inserted into a membrane composed of 1-palmitoyl-2-oleoyl-sn-glycero-3-phosphatidylcholine (POPC) and cholesterol (CHL) in a 7:3 ratio, using CHARMM-GUI bilayer membrane builder.^62,63^ The protein’s orientation in the bilayer was obtained from the Orientation of Protein and Membranes (OPM) database.^64^ A 22 Å-thick layer of water was added above and below the membrane along the z-axis, and NaCl ions were used to neutralize the system while maintaining a salt concentration of 0.15 M. All components considered, our systems have, on average, 150,000 atoms in total for the inactive receptor states, while the miniG-bound ARs have up to 175,000 atoms. Simulated systems were set up using the *tleap* utility in *AmberTools22* using the Amber 14SB force field for the protein, Lipid17 for the membrane, and TIP3P for water and Joung-Chetham parameters for the monovalent ions.^65–69^ The ligands’ force field parameters were generated by performing geometry and charge optimizations using the Gaussian16 implementation of the Hartree-Fock level of theory with a 6-31+G** basis set.^70,71^ We then assigned the partial charges to the atoms using RESP, and the angle, dihedral, and bond parameters were assigned by Amber’s Gaff2, utilizing the antechamber tool.^72–74^ The energy minimization, equilibration, and production run simulations were performed using GROMACS 2020 patched with PLUMED 2.7.^75,76^ After an energy minimization step to remove possible steric clashes introduced by the CHARMM-GUI bilayer builder, the systems were thermalized using 9 cycles of NPT and NVT steps and a pre-production run of 20 ns. The subsequent production run simulations were performed under the NPT ensemble with a V-rescale thermostat and a Parrinello-Rahman barostat.^77,78^

### Thermodynamic calculations (Funnel-Metadynamics)

For the FM calculations, we placed a funnel-shaped restraint potential covering the OBS of the ARs, using the VMD tool *funnel.tcl* developed by Raniolo *et al.*^79^ To sample a comprehensive phase space of the ligand-bound states, we used two geometric collective variables (CVs) throughout all of our ligand-protein complexes. These CVs are (1) the distance between the center of mass (COM) of the ligand and the COM of the OBS of the protein as defined by the *C_α_* positions of selected residues and (2) a torsion angle that considers the rotation of the ligand projected into a plane that is parallel to the plane of the base-like moiety of the adenosine receptor ligands when bound. See the supplementary section “CV Explanation” for figures and further discussion on how these CVs are defined. The *σ* values for the well-tempered metadynamics Gaussian-shaped potential acting on the biased CVs are taken in proportion to the standard deviation of these values when measured in unbiased MD simulations performed for each system. The bias potential is deposited every 500 steps (i.e., 1 ps) with a bias factor equal to 20 and an initial height equal to 2.0 kJ/mol.

### Kinetics calculations (Infrequent Metadynamics)

For the AdInf simulations, we perform a large number of short parallel and independent simulations, each with a maximum length of 10 ns, starting from the most stable binding pose as determined by our FM approach. We accelerated the unbinding pathway sampling by adding a bias on a protein–ligand distance CV at a variable deposition rate with a minimum rate of 1 Gaussian deposition every 10 ps. The simulation stops when the ligand reaches the unbound state using the PLUMED analysis module COMMITTOR. We use the unbinding data to build a state-to-state Markov transition matrix that describes the ligand’s transition probability and residence times (RT) between the main binding poses and intermediate states. We validate the kinetics statistics by performing a KS test with a theoretical exponential distribution as proposed by Salvalaglio *et al.*^44^ In this KS test, we test the null hypothesis by comparing our empirical exponential cumulative distribution function (ECDF) of residence times to a theoretical exponential cumulative distribution function (TCDF). We fit the two CDFs by using the Python *SciPy* library *curve fit* function.^80^ We then visualize and analyze the unbinding trajectories by mapping the trajectories into the same CV space as used for the thermodynamic calculations (i.e., distance and torsion angle). In the CV space, we can then create unbinding heatmaps to illustrate the most visited metastable states for the unbinding pathway(s). We create these heatmaps using Python’s *NumPy* function *histogram2d*, onto which we apply a Python *SciPy* gaussian filter as a kernel to create a smooth histogram.^80,81^ We use the heatmap hotspots to identify and define the metastable states of the transition matrix M. Each entry of the matrix M*_ij_* contains the information on the RT for the ligand as it transitions from state *i* to state *j* and the probability of this transition, as was similarly done in Tiwary *et al.* but without the need to perform a new AdInf from each of the defined basins.^43^ From the matrix M, we can easily identify which is the rate-determining transition for each ligand and map each of these CV states into their structural representation. We then explore the CV space along this transition to identify the transition-state (TS) configuration. The TS is validated through 100 independent simulations initiated from this configuration, confirming that approximately 50% of the trajectories reach one basin and 50% reach the other. Please refer to the Supplementary Information section *Kinetics* to see the heatmaps and the complete transition matrix M for each ligand-protein pair. The extraction of the structures in the representative metastable states is detailed in the Supplementary Information section *Representative metastable states*.

## Results and Discussion

In the following section, we present the thermodynamic and kinetic data for TEP, ZMA, and NECA in complex with A2A and A2B receptors.

### Theophylline - the antagonist

#### Thermodynamics

Theophylline (TEP), also known as 1,3-dimethylxanthine, is a non-selective A2A/A2B antagonist.^82^ In the case of A2A, TEP’s most stable binding mode has been experimentally elucidated in two different studies.^48,83^ Conversely, no experimental structure of the TEP-A2B complex has been resolved to date. In this section, we present and discuss the most energetically relevant binding modes we compute for TEP on both receptors using the FM protocol. We highlight the main similarities and differences that TEP exhibits while bound to both receptors, with the scope of rationalizing their relative affinity. In Fig. 3, we present the thermodynamic and kinetic profiles of TEP binding in A2A and A2B. By comparing the FES of both ARs as a function of two biased CVs, i.e., ligand-protein distance and torsion angle, we observed qualitatively similar profiles (Fig. 3a, b). Overall, the most notable difference lies in the distribution of low-energy-bound states (i.e., states with a distance CV less than 1 nm), which show greater degrees of diffusion along the torsional degree axis (y-axis) in A2B than in A2A. This suggests that TEP exhibits greater torsional degrees of freedom within the A2B’s OBS, likely due to subtle structural differences between the two receptors, such as the mutation L^6.51^ in A2A to V^6.51^ in A2B. Despite these differences, the global minimum energy state for both receptors (labeled as state A in Fig. 3, **a** and **b**) is remarkably similar, with an RMSD of 0.95 Å of TEP’s heavy atoms between A2A and A2B for the binding pose A. This finding is consistent with the available experimental evidence for the A2A-TEP experimental structure, for which we obtain an RMSD of 0.2 Å between A2A’s pose A and the crystal binding pose. Thus, our results demonstrate that the lowest-energy binding mode of theophylline in the OBS is conserved between the A2A and A2B receptors. From a thermodynamic perspective, we computed binding free energy (ΔG) of −6.89 ± 0.14 kcal/mol for A2A and –6.74 ± 0.31 kcal/mol for A2B, in agreement with experimental measurements that fall within the ranges of [–7.08; –7.90] kcal/mol for A2A and [–5.58; –6.81] kcal/mol for A2B, reproducing the non-selective binding profile of TEP.^84–86^ The most important residue interactions that stabilize state A in A2A and A2B are a *π* − *π* stack with F168(A2A)/F173(A2B), and a strong double H-bond with N^6.55^, which is in agreement with experimental observations reported by Carpenter *et al.*^48,49,87^ Thus, the computation of similar low-energy binding modes aligns well with the expected behavior given the structural similarity of the OBS of the two receptors. Interestingly, we identified a second low-energy minimum in A2B (labeled as state B in Fig. 3 **b**), which is found in equilibrium with state A. Please refer to the Supplementary Section *Alternative Binding Modes* to read the discussion on this pose.

**Figure 3:**
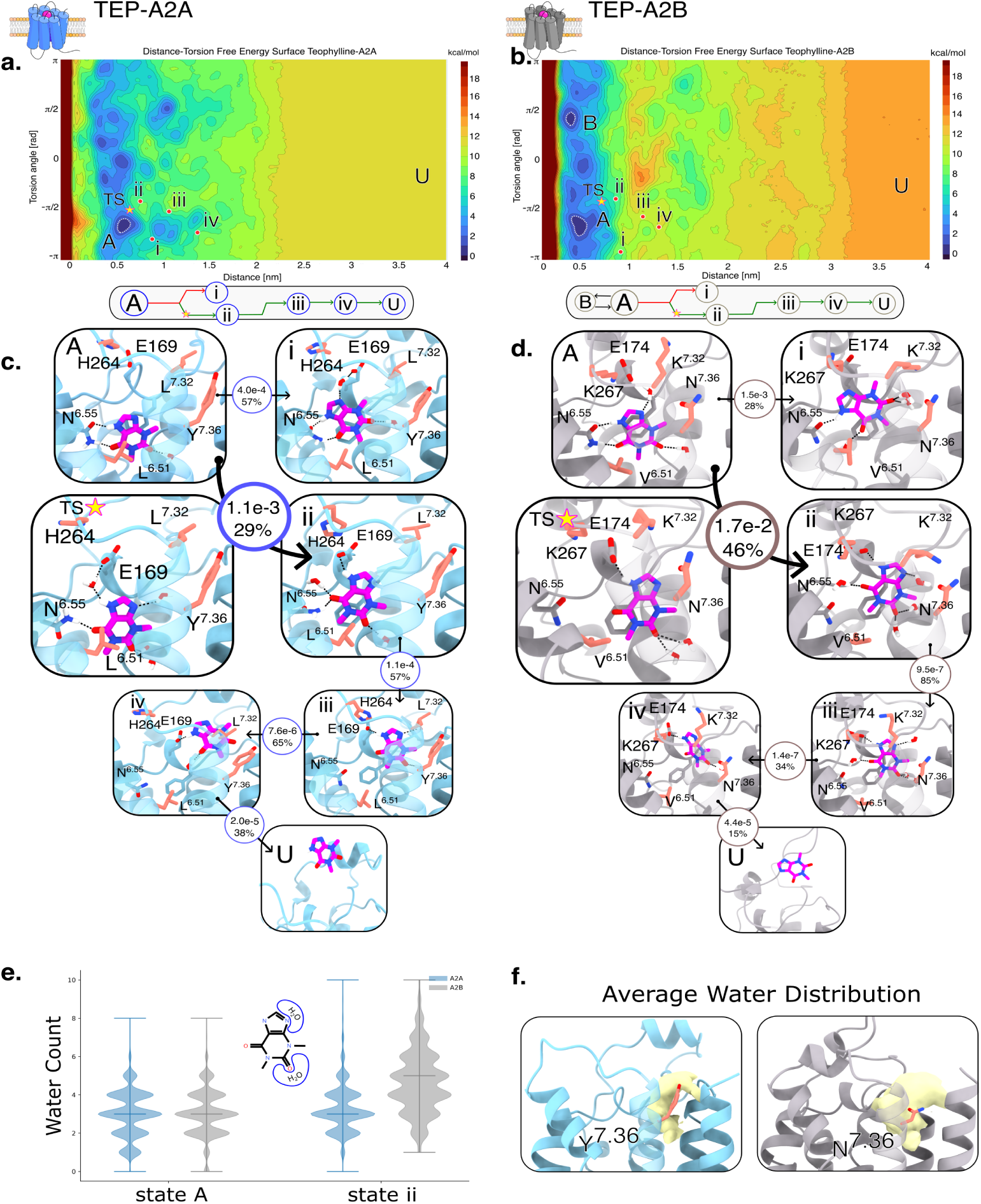
**a** Shows the contour plot of the computed FES as a function of the two CVs (distance and torsion) for the TEP-A2A complex. **b** Shows the same computation but for the TEP-A2B complex. In both plots, the contour isolines are shown every 1 kcal/mol. The label A in each figure indicates the CV coordinate of the global energetic minimum. The label B indicates the coordinates of the second-lowest global minimum. The orange star labeled TS indicates the CV coordinates of the transition state for the most relevant unbinding pathway. The lower-case Roman numerals indicate the CV coordinates of the kinetic metastable states where TEP resides for longer times while unbinding from the protein. The U label indicates the high isoenergetic region unbound state for the complex. Below each contour plot, we show a transition diagram that indicates the main flux that TEP exhibits with the receptors. Immediately beneath, in panel **c**, we show the structures that represent each of the aforementioned labels. For A2A, the protein is shown in light blue, while for A2B, the protein is shown in light gray. In both figures, TEP is shown in magenta. The protein residues that are the same for both receptors are shown in their native protein color, while point mutations in the orthosteric binding site are colored in salmon. Each structure shown is labeled with either letters or Roman numerals, which map directly to its related CV coordinate as shown in the FES. Furthermore, we show the RT of each state along with its associated probability of transition. The only unlabeled residue is *F* 168(A2A)/*F* 172(A2B) to promote image clarity. Panel **e** shows the distribution of water molecules that come within 4 Å of theophylline atoms N9 and O11 in state A and state *ii*. Panel **f** shows the average spatial distribution of water molec1u3les around the residues *Y/N* ^7.36^.

#### Kinetics

Despite the structural similarities between the OBSs of A2A and A2B, the computed RTs of TEP differ by one order of magnitude between the two receptor subtypes, with values of 1.62e-3 s ± 2.1e-4 s for A2A and 2.56e-2 s ± 3.5e-3 s for A2B. To our knowledge, only one study has reported an *in vitro* RT for TEP at A2A, yielding a value of approximately 13 min at 10*^◦^*C, rendering a direct comparison difficult.^60^ For A2B, we did not identify any *in vitro* kinetic measurement against which our estimate could be compared. Nevertheless, the relative difference of our calculated RT between the two receptor subtypes remains informative. In the following section, we rationalize the approximately one-order-of-magnitude difference in the calculated RTs in terms of the distinct unbinding mechanisms and receptor-specific structural features observed for A2A and A2B.

Starting from the lowest-energy bound state (state A) identified in our thermodynamic analysis, we observed four main metastable states that lead to the unbinding event of TEP from A2A and A2B. For clarity, these identified metastable states are mapped (using lowercase Roman numerals) in the FES derived from our thermodynamic simulations (Fig. 3 **a**, **b**), whilst the rate-determining TS are marked with a star on the same figure. Furthermore, below each FES, we have created a transition diagram that helps understand the main unbinding transitions that the ligand executes while leaving the receptors. In the A2A receptor, the main unbinding pathway of TEP follows a transition from state A to state *ii* with a 29% probability and an RT of 1.0e-3 s. The TS of TEP in A2A is characterized by a slight increase in its torsion angle (clockwise rotation) relative to state A. This reorientation is facilitated by the interposition of a water molecule between the N7-hydrogen of TEP and the nitrogen-bound hydrogen in the side chain of N^6.55^ and the formation of an H-bond between TEP’s O13 and N^6.55^’s nitrogen. Moreover, the water molecule that inserts itself between TEP and N^6.55^ performs a water-mediated bridge between the ligand and E169. (Fig. 3 **c**). This water-mediated interaction then promotes the ligand to commit to state *ii*, where it interacts directly with E169 and N^6.55^. From state *ii*, with an RT of 1.4e-4 s, it transitions into state *iii*, where the ligand continues to interact directly with E169 but loses direct contact with N^6.55^ and with F168. At this stage, TEP is located in what we call a hydration gate formed by the space between TM1, TM2, and TM7, and modulated by Y^7.36^(N in A2B). See, for example, that TEP’s O11 is in contact with a water molecule in front of Y^7.36^(N in A2B) and TEP’s N9 with water molecules in front of L^7.32^(K in A2B). Afterwards, with an RT of 7.6e-6 s, TEP continues to transit to state *iv* by moving upwards in the pocket, keeping direct contact with E169 and forming a direct VdW contact with L^7.32^(K in A2B). At this state, the ligand is at the vestibule between the orthosteric pocket and the solvent state, where it transits in 2.0e-5 s to the unbound solvent state. For completeness, starting from state A, TEP also transitions to state *i*, with lower RT and higher probability than to state *ii*. In this state, TEP maintains strong direct interactions with N^6.55^ and water-mediated interactions with E169. However, this pathway does not lead to productive unbinding because its upward movement is impeded by the salt bridge formed between H264(K in A2B) and E169. This salt blockage essentially pushes the ligand back into state A, rendering the main transition pathway from state A to state *ii* the unique most relevant unbinding path. Interestingly, the unbinding mechanism of TEP from the A2B receptor exhibits similar processes to the ones observed in A2A (Fig. 3 **d**). Specifically, its metastable state *ii* is strikingly similar to the one observed in A2A. However, the TS that leads to this transition is different. The TS in A2B does not present an insertion of a water molecule between the TEP and N^6.55^, but instead we observe a spontaneous loss of H-bond interactions with N^6.55^ and a formation of an H-bond with E174, which then leads to state*ii* with an RT of 1.6e-2 s. Once in state *ii*, TEP is stabilized by a water-mediated H-bond with N^6.55^ and by the direct H-bond with E174, which is extremely similar to state *ii* in A2A except for the direct contact between TEP and N^6.55^. We note that in this state, TEP is experiencing a larger number of water contacts via its O11 and N9 than when compared with its A2A counterpart. Relatively quickly, TEP transitions to state *iii*, where it loses direct interactions with F172, maintains a water-mediated H-bond with N^6.55^, and keeps the direct H-bond with E174; it also enters into a network of water-mediated interactions held by N^7.36^. Here, TEP inserts itself into what we previously denominated as the hydration gate formed by TM1, TM2, and TM7, but as opposed to A2A, in A2B, this gate is modulated by N^7.36^. We hypothesize that the presence of N^7.36^ in A2B, as opposed to Y^7.36^ in A2A, increases local hydration between TM1, TM2 and TM7. From state *iii*, TEP quickly transitions to the pre-unbound state *iv*, where the interaction with the key residue N^6.55^ is entirely lost, a direct H-bond with E174 is still maintained, and the ligand is right in front of K^7.32^(L in A2A). Afterwards, TEP completely unbinds from the pocket and loses all interactions with the receptor. Alternatively, and similar to what occurs in A2A, state A can also evolve towards state *i*. In this conformation, a water molecule interposes between N^7.36^ and the ligand, forcing TEP to rotate approximately 45 degrees in the counterclockwise orientation relative to state A. But again, this state rarely leads to a successful unbinding event because its outward movement from that conformation is blocked by the salt bridge E174 - K267(H in A2A).

We believe that the similarity in the main unbinding pathway characteristics between the two receptors may be attributed to the salt bridge between E169 - H264 in A2A and E174 - K267 in A2B, which partially occlude the vestibular lumen, constraining the ligand to orient itself towards TM1 to egress from the OBS. Yet they present different RT and transition probabilities, and we believe that these differences are driven by the number of water molecules that come into contact with the ligand, especially in the first metastable states, such as in state *ii*. We further investigated this phenomenon and discovered that, indeed, TEP, after going through the TS state, experiences two distinct environments on the two receptors. In Figure 3, panel **e**, we see that the distribution of water molecules that come within 4 Å of the atoms O11 and N9 is different between state A and state *ii* for A2A and A2B. In state A, TEP experiences similar levels of hydration on A2A and A2B; but when the ligand is found in state *ii*, interestingly, the hydration level stays mostly the same for A2A when compared to state A, but for A2B, the hydration contacts notably increase. This increase in water contacts is visually seen in the panels **c** and **d** in Figure 3, where we see that TEP in state *ii* on A2A has one direct H-bond water contact through the O11 atom, whilst for A2B, TEP exhibits three water molecule contacts, two with O11 and one with N9. As previously introduced, we believe that the mutation Y/N^7.36^ creates a wholly different water environment in this region of the receptor. We quantified this difference by generating a Volumetric Density Map (VolMap) of the water molecules that are around these residues and show it in Figure 3, panel **f**. The yellow surface represents the average presence of water molecules around these residues throughout a 1 *µ*s unbiased MD simulation of each receptor in complex with TEP. On A2A, we can visually appreciate that there is a crescent moon shape that goes around Y^7.36^. The surface shape on A2B is more continuous, as it is less correlated to the position of N^7.36^. We quantified the number of water molecules that define this VolMap and plot the distribution in the Supplementary section *Y/N*^7.36^ *Hydration Gate*. The distributions show that in A2A there are, on average, 7 water molecules in this region, while in A2B there are about 11 water molecules; furthermore, the distributions show totally different minimum and maximum values, further supporting the idea that A2A’s water presence in this hydration gate is notably lower than for A2B. The conclusion is that this Y/N^7.36^ mutation does indeed contribute to the water molecule presence and diffusion into the receptors OBS. This region, comprised of TM1, TM2, and TM7, has been discussed previously by Chen *et al.*, who call it a secondary pocket in which the resolved binding pose of the agonist BAY60-6583 directly interacts with N^7.36^ with its polar tail.^24^ This ligand is highly selective for A2B, and when the authors mutated N^7.36^ to alanine, the ligand suffered a 6-fold decrease in potency.^24,88^ Conversely, in A2A, the residue Y^7.36^(N in A2B) has been identified as a key residue that, when mutated to alanine, completely disrupts the binding of the inverse agonist ZMA, the antagonist XAC, and the agonist CGS21680.^89,90^ Certainly, the polarity introduced by N^7.36^ in A2B and the hydrophobicity introduced by Y^7.36^ in A2A, whether through direct ligand contacts or through their effects on local hydration, represent not only thermodynamic but, as we have shown, also kinetic determinants.

### ZM-241385 - the inverse agonist

#### Thermodynamics

In the following sections, we report the thermodynamic and kinetic calculations for the A2A-selective inverse agonist ZMA in complex with A2A and A2B.^47^ The ZMA-A2A complex has been extensively studied, with over 30 experimentally resolved structures published.^91^ On the contrary, the ZMA-A2B complex structure has not been resolved to date. It has been shown, however, that ZMA binds to both receptors, albeit with a higher affinity for A2A.^92,93^ In Fig. 4, **a** and **b**, we show the FES computed through FM as a function of the two biased CVs (distance and torsion) for ZMA in complex with A2A and A2B, respectively. The FES profiles for both receptors are quite similar; both exhibit the existence of a single binding mode, illustrated by having only one significant minimum. This is expected due to the steric hindrance of ZMA, whose elongated shape appears to favor only one preferential conformation in the OBS of both receptor subtypes. The absolute binding free energy ΔG computed for the ZMA-A2A complex is −11.01 ± 0.86 kcal/mol, close to the reported experimental values of [−11.9 to −12.4] kcal/mol.^92^ For the ZMA-A2B complex, the computed ΔG is −8.28 ± 0.28 kcal/mol, in line with the reported experimental values of [−9.2 to −11.17].^93–95^ Below both FES figures, we show the binding mode that represents the poses that define the lowest-energy binding mode (state A) (Fig. 4c). The identified binding mode shown in Fig. 4 panel **a** is strikingly similar to the experimental binding pose (PDB: 5IU4) with an RMSD of 0.86 Å, computed over all heavy atoms.^87^ The ZMA binding mode in the A2B receptor (Fig. 4 panel **b**) closely matches the binding mode of ZMA in A2A, with an RMSD value of 1.66 Å between them. Indeed, analysis of the most stable binding modes of the two compounds reveals that the interaction patterns established between the ligand and the receptor are mostly conserved across A2A and A2B subtypes. Given the resemblance of the two binding modes and the conservation of the interacting residues, it is interesting to take a closer look at the structural determinants underlying ZMA selectivity towards the A2A receptor. Notably, both receptors display a conserved double H-bond with N^6.55^ and a hydrophobic *π* − *π* stacking interaction with F168(A2A)/F172(A2B), as is common for all adenosine receptor binders. The flexible glutamate located in the ECL2 region (E169 in A2A and E174 in A2B) further stabilizes ZMA by coordinating its free amine group, either via water-mediated interaction or via direct contact. The main structural difference that comes into direct contact with ZMA at the OBS level involves residue L^6.51^ in A2A, which is mutated to V^6.51^ in A2B.^24^ Thermodynamically, this residue L/V^6.51^ appears to constitute a key spot for the selective recognition of A2A’s inverse agonist ZMA, which is consistent with previous reports.^96^ Since this conservative amino acid substitution does not alter the polarity of the OBS floor, we hypothesize that the effect of this spot on ZMA selectivity may be attributed, at least in part, to the distinct steric hindrance between valine and the bulkier side chain of leucine. To test this hypothesis, we quantified the volume of the OBS floor during a 1 *µ*s unbiased MD simulation of both A2A and A2B receptors in complex with ZMA. Overall, the OBS floor volume was, on average, 15 Å^3^ larger in A2B than in A2A. The two systems also exhibited distinct volume distributions, particularly toward the lower end of the sampled range. While A2A reached minimum OBS volumes as low as 49 Å^3^, the minimum observed for A2B was 77 Å^3^, indicating that A2B consistently maintained a larger OBS floor volume (see Supplementary Information, section *OBS Volumes*). As a consequence, ZMA in A2A is stabilized through a stronger VdW packing of L^6.51^(V in A2B) against the bicyclic triazolotriazine-furan core of the ligand. We hypothesize that this interaction promotes a tighter insertion of the furan moiety into the reduced OBS volume of A2A, in a manner more favorable than within the larger cavity of A2B. To test this hypothesis, we monitored the torsion angle CV during unbiased MD of minima to assess the torsional stability of the ligand. We measured higher degrees of freedom of ZMA in A2B’s OBS, which could be directly correlated to its larger volume (See Supplementary Information section *Unbiased MD Simulations of Thermodynamic Minima*). This behavior likely arises from the loss of structural constraints imposed by the L^6.51^(V in A2B) residue, which is no longer maintained in A2B. Consistently, the RMSD of ZMA’s heavy atoms during unbiased simulations appears to be significantly lower and more stable in the A2A system when compared to ZMA in A2B, further supporting this observation (see Supplementary Information section *Unbiased MD Simulations of Thermodynamic Minima*). These observations suggest that the preference for A2A may not be restricted to ligands containing the exact triazolotriazine-furan scaffold of ZMA, but could extend more broadly to antagonists sharing a similar structural motif, namely a fused bicyclic nitrogen-containing heteroaromatic core connected to a furan ring through a single bond and adopting a comparable binding orientation. Under the assumption that these ligands engage the two receptors in a similar manner, such a scaffold may favor binding to A2A over A2B. Consistent with this hypothesis, antagonist ligands in the Guide to Pharmacology database displaying this general structural arrangement show higher reported affinity for A2A than for A2B. Examples include SCH442416, preladenant, and SCH58261, which retain a closely related bicyclic heteroaromatic-furan architecture despite differences in the precise chemical identity of the fused bicyclic core.

**Figure 4:**
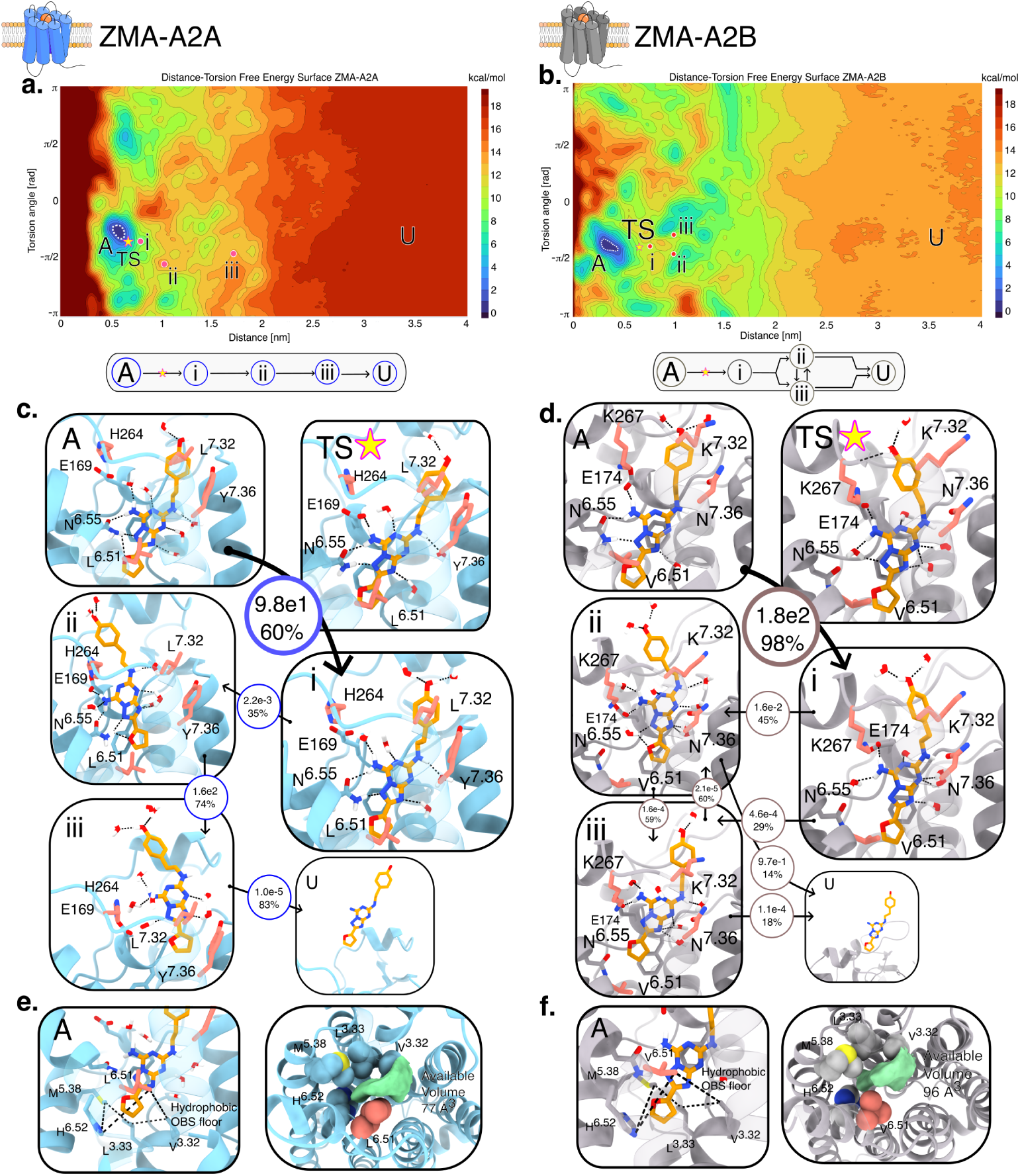
**a** Shows the contour plot of the computed FES as a function of the two CVs (distance and torsion) for the ZMA-A2A complex. **b** Shows the same computation but for the ZMA-A2B complex. In both plots, the contour isolines are shown every 1 kcal/mol. The label A in each figure indicates the CV coordinate of the global energetic minimum. The label TS indicates the coordinate of the transition state for the most relevant unbinding pathway. The lowercase Roman numerals indicate the coordinates of the metastable states, i.e., where ZMA resides for longer times while unbinding from the protein. The U label indicates the high isoenergetic region for the unbound state of the complex. Below each contour plot, we show a transition diagram that indicates the transitions that ZMA exhibits with the receptors. Immediately beneath, we show the structures that represent each of the aforementioned labels. For A2A, the protein is shown in light blue, while for A2B, the protein is shown in light gray. In both figures, ZMA is shown in orange. The protein residues that are the same for both receptors are shown in their native protein color, while point mutations in the orthosteric binding site are colored in salmon. **c** shows the most probable transitions for the unbinding of ZMA from A2A. **d** shows the most probable transitions for the unbinding of ZMA from A2B. Each structure shown is labeled with either letters or Roman numerals, which map directly to its related CV coordinate as shown in the FES. Furthermore, we show the RT of each state along with its associated probability of transition. The only unlabeled residue is F168(A2A)/F172(A2B) to promote image clarity. **e** and **f** show the hydrophobic floor subpocket formed by the residues V^3.32^, L^3.33^, M^5.38^, and H^6.52^ on both receptors. The furan moiety of ZMA is inserted into this tight VdW pocket. Furthermore, in these panels we show a top view of the receptors to highlight the volume of the cavity formed by these residues. We make note of how much larger the available space is in A2B when compared to A2A, visually and numerically, with these surfaces measuring 93 and 77 Å^3^ for A2B and A2A, respectively.

#### Kinetics

The unbinding of ZMA from A2A and A2B shares some common features, but the overall pathways are largely distinct. In this section, we describe the characteristic structural features and RTs associated with the most favorable unbinding pathway identified for ZMA in each receptor using the AdInf approach described above (see Methods). We estimate an RT of 302 s ± 40 s for ZMA bound to A2A and 180 s ± 23 s for ZMA bound to A2B. Several *in vitro* studies have reported ZMA binding kinetics at A2A under different experimental conditions and temperatures.^60,89,97–101^ Among these, the studies by Bocquet *et al.* and Pettersen *et al.* employed experimental systems that most closely resemble our MD setup.^97,98^ These studies reported A2A residence times for ZMA of approximately 280-320 s and 19 s, respectively. Our calculated RT is therefore in strong agreement with the values reported by Bocquet *et al.* To our knowledge, no *in vitro* kinetic measurements of ZMA binding to A2B have been reported to date. In the following section, we rationalize the difference in our calculated RTs between A2A and A2B.

In A2A, the initial energetic barrier for ZMA egress (labeled as TS) is very close to the most stable binding mode when measured in the distance-torsion CV space (see Fig.4 panel **a**). From our determination of the TS, we observed that the unbinding of the furan group, which is buried in the hydrophobic OBS floor subpocket formed by V^3.32^, L^3.33^, M^5.38^, L^6.51^(V in A2B), and H^6.52^ (see 4 panels **e** and **f**), presents the highest energetic barrier. This hydrophobic pocket, aided by the point mutation L^6.51^(V in A2B), creates a tight space for ZMA to unravel itself from, and thus energetically disfavors the unbinding of ZMA. In comparison, in state A (Fig. 4 panel **c**), the furan moiety is deeply buried within the pocket, where L^6.51^(V in A2B) is essentially above it. In contrast, in the TS, the ligand has moved upwards, and the furan ring is collinear with L^6.51^(V in A2B), which then motivates the system to relax at a higher distance, as shown in state *i*. At this stage, the furan moiety is outside A2A’s hydrophobic pocket, and although the important contact with N^6.55^ is maintained, the *π*-*π* stack with F168 is no longer optimal. Overcoming these volume and chemical challenges renders state A with an RT of 98 s. Afterwards, ZMA quickly transits to state *ii*. The contacts formed in this state are practically identical to state A, with the exception that the *π*-*π* stack with F168 is almost lost and the furan moiety has completely lost all interactions with the hydrophobic pocket. This state is thermodynamically stable, and it presents a high RT of 160s before transitioning to state *iii*. State *iii* is a complete loss of all OBS contacts, and ZMA is maintained by the vestibular contacts such as the point mutations L^7.32^(K in A2B), Y^7.36^(N in A2B), and an array of water networks. Being already in a pre-unbound state, ZMA then quickly transits to the fully unbound states.

As previously discussed, the bound state A of ZMA in A2B is very similar to the bound state A of A2A. The unbinding kinetics, however, are noticeably different. First, ZMA presents a single high-energy unbinding stage that is responsible for the overall unbinding RT, that is, the transition from state A to state *i*. The process it goes through is of a similar nature to that in A2A; that is, the first step of ZMA is to leave the hydrophobic pocket on the OBS floor. However, in A2B, the transition from state A to state *i* involves the occurrence of other factors that are not seen on the TS of A2A. Namely, not only does the furan moiety leave the hydrophobic pocket, but state *i* also does not present any direct contact with N^6.55^, and instead it interacts with it through a water bridge. Moreover, as seen in the case of TEP-A2B, this state presents a notably higher degree of water molecules in contact with ZMA. As analyzed in the thermodynamic section, the volume of the hydrophobic subpocket formed by V^3.32^, L^3.33^, M^5.38^, L^6.51^(V in A2B), and H^6.52^ is larger in A2B; thus, we expect that ZMA does ‘slip’ out faster from this loosely fitting pocket than for A2A. However, the RT to leave this hydrophobic pocket seems to be longer in A2B. We attribute the longer RT in A2B to the fact that ZMA, to reach state *i*, has to coordinate many of the factors just described, which render state A with an RT of 180 s, higher than the comparable counterpart of A2A. State *i* however, keeps direct contact with E174. Afterwards, ZMA’s pathway bifurcates into state *ii* or *iii*. Both of which place ZMA closer to a pre-unbound state where the *π* − *π* interaction with F172 is lost, and both maintain a direct H-bond with E174. The difference between these two states is a few degrees in rotation from the torsion angle CV point of view; thus, these two states are in a sort of rapid equilibrium. Both of these states then contribute almost equally to the following unbinding step, albeit with slightly different RT.

### NECA - the agonist

#### Thermodynamics

In the case of the non-selective agonist NECA, solved structures are available for both receptors (PDB IDs: 2YDO, 5G53 for A2A, and 7XY7 for A2B).^24,49,102^ The 2YDO structure resolved NECA in a complex with an intermediate active state of A2A. Meanwhile, the 5G53 and 7XY7 structures resolved NECA in the fully activated states of the receptors, coupled with an engineered miniG*_s_* in the case of A2A and with the complete heterotrimeric G-protein effector in the case of A2B. As detailed in the Methods section, we study NECA in complex with the fully activated states of the receptors, i.e., A2A and A2B are both coupled with the engineered miniG*_s_*-protein throughout the thermodynamic and kinetic studies. In Fig. 5, panels **a** and **b**, we show the main results of the thermodynamic studies for the A2A and A2B systems, respectively. The FES between the two receptors is topologically different. The A2A system shows a global minimum labeled as A and then a significantly low-energy state which coincides with the CV coordinates of the kinetically metastable state labeled as *ii* (see the following kinetics section for details). Meanwhile, the A2B system exhibits only one stable bound pose with a region of high energy values that seems to separate the bound from the unbound states, a feature that is not present in A2A’s FES. Below both FES figures, we show a representative binding pose that populates the global minimum (state A) along with all the residues that form strong interactions with the ligand. These include: T^3.36^, E169/E174, H^6.52^, N^6.55^, H^7.43^, and S^7.42^, and the *π*-*π* stack with F168/F173. The binding poses obtained for both receptors are extremely similar to their respective experimentally resolved binding poses, with an RMSD of 0.44 Å in A2A and 0.55 Å in A2B for all heavy atoms when compared with the poses reported in PDB IDs 5G53 and 7XY7, respectively.^24,102^ In contrast to TEP and ZMA, NECA performs agonist-specific contacts with residues located on TM3 and TM7, such as T^3.36^, H^7.43^, and S^7.42^, as reported by Carpenter *et al.* (although there has been recent evidence of inverse agonists interacting with A2A’s T^3.36^).^87,103^ We compute a ΔG of −9.05 ± 0.5 kcal/mol for NECA in A2A, in agreement with the experimental range of [−9.40 to −11.85] kcal/mol.^88,104^ While for A2B we compute a ΔG of −8.98 ± 0.12 kcal/mol, that falls in the range of reported experimental values [−7.78 to −9.40] kcal/mol.^88,104^

**Figure 5:**
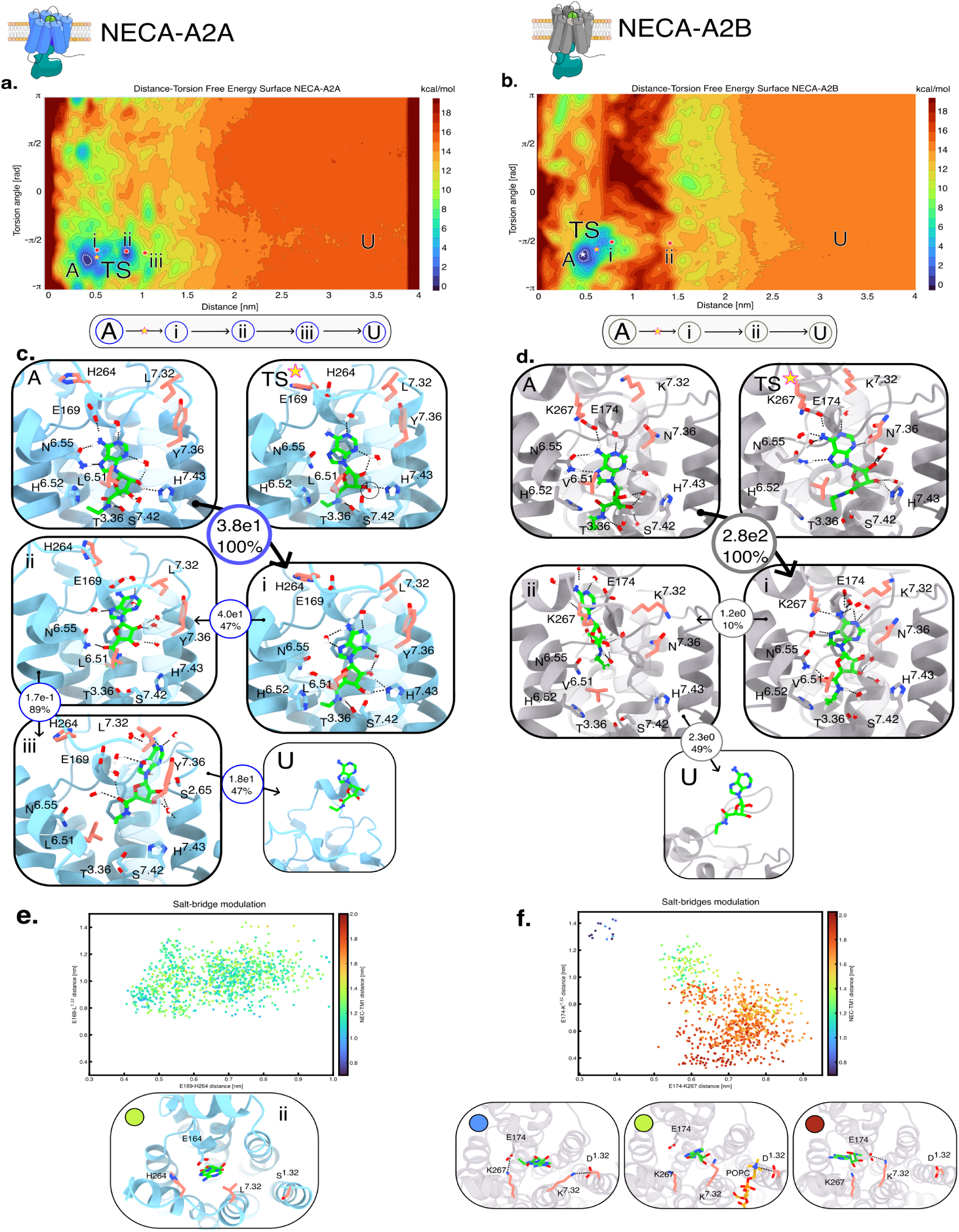
**a** Shows the contour plot of the computed FES as a function of the two CVs (distance and torsion) for the NECA-A2A complex. **b** Shows the same computation but for the NECA-A2B complex. In both plots, the contour isolines are shown at intervals of 1 kcal/mol. The label A in each figure indicates the CV coordinate of the global energetic minimum. The label TS indicates the coordinate of the transition state for the most relevant unbinding pathway. The lowercase Roman numerals indicate the coordinates where NECA resides for longer times while unbinding from the protein. The U label indicates that the high isoenergetic region is the unbound state for the complex. **c** Below each contour plot, we show the structure that represents the global minimum A, along with the most probable transitions for the unbinding of NECA from A2A. **d** shows the most probable transitions for the unbinding of NECA from A2B. Each structure shown is labeled with either letters or Roman numerals, which map directly to its related CV coordinate as shown in the FES. Furthermore, we show the RT of each state along with its associated probability of transition. **e** shows a scatter plot of the distance between the salt bridge E169-H264 and E169-L^7.32^, while **f** shows a scatter plot of the distance between the salt bridges E174-K267 and E174-K^7.32^. Both scatter plots are colored by the distance between NECA and the top part of TM1. The scatter plots show that NECA, state *ii* on A2B, has two exit pathways, with three protein conditions, colored by blue, green, and red. While the same analysis shows that NECA *ii* exhibits only one pathway.

#### Kinetics

The kinetics for the unbinding of NECA are notably different between the two receptors. The RT computed for the unbinding time of NECA from A2A is 72 s ± 11 s, whereas for A2B it is 290 s ± 50 s. We are able to compare our both of our *in silico* estimates with various *in vitro* experiments.^98–100,105^ From these studies, we remark that our estimate for A2A-NECA is remarkably close to the assay performed by Pettersen *et al*, in which they calculate an RT of 25 s. Whilst for A2B, the only *in vitro* estimate we found to compare with reports is an estimate of 35 s.^105^

Interestingly, NECA in A2A seems to exhibit a two-high RT step process, while in A2B, it exhibits a single high RT step during its unbinding process. In Fig.5, panels **c** and **d**, we report the metastable states of NECA while it unbinds from A2A and A2B, respectively. The transition from state A to state *i* in A2A requires an increased rotation of the purine moiety in the clockwise direction, increasing its distance from the key residue N^6.55^ and allowing the insertion of a water molecule in between. This transition is facilitated by an increase in solvation of the OBS pocket and a stochastic loss of direct interaction with E169. This is exemplified by the TS state, where water molecules get inserted between NECA and the agonist-specific residues S^7.42^ and H^7.43^, thus destabilizing the ligand’s ribose moiety, and E169 moves up, increasing the available space for NECA to perform its rotation. One such water molecule insertion is circled in Figure 5, panel **c**, TS state. This then promotes the ligand to higher energy states, reaching the metastable state *i*. Therefore, state *i* is characterized by the loss of direct contact with N^6.55^ and E169, and by an increase in water-mediated contacts between NECA and the protein. This endows state A with an RT of 38 s. However, state *i* is still a relatively stable bound state. This is due to the fact that the ribose moiety, along with the 5’-N-ethylcarboxamide group, is in a position that is highly similar to their position in state A. This means that the majority of the stabilizing contacts that endow state A with its low free energy are still present for these parts of NECA. Then, state *i* transitions to state *ii*, with a comparable RT of 40 s, at which NECA is approaching the vestibular binding site of the protein, effectively losing almost all the contacts found in state A. State *ii* is characterized by a disruption of the E169-H264(K in A2B) salt bridge, a higher number of water contacts provided by the hydration gate at Y^7.36^, loss of the optimized *π*-*π* stack with F168, and the formation of a direct H-bond between O5’ and N^6.55^. State *ii* is a high-energy state with a low RT, where NECA transitions to the pre-unbound state *iii*, where it is mostly kept in that position via a complex network of water-molecule bridges and an H-bond with S^2.65^. The next step is a complete unbinding of the ligand from the protein. In the case of A2B, NECA’s first transition is very similar to the one in A2A, with some defining differences between them. Firstly, NECA’s state A is a more hydrated state when compared to A2A’s state A. We note this by seeing the water molecules that are lined up and form a column of hydration below N^7.36^ to H^7.43^. Already here, we notice a striking difference between NECA in the two receptors. Whilst in A2A, NECA’s TS is determined by the insertion of water molecules between H^7.43^ and the ligand; in A2B, these water molecules are already present in A2B’s state A. Although state *i* is somewhat similar between the receptors, the way each gets there is totally different. In A2B, the key rotation is given by the ribose moiety, as opposed to the base in A2A, where contacts with T^3.36^ and S^7.42^ are lost, but direct contact with H^7.43^ is formed. Moreover, water molecules interrupt the optimized double H-bond with N^6.55^, but crucially, NECA maintains a direct contact with E169. This movement is spatially larger than the one NECA performs in A2A, and it is presumably a larger leap in free energy. This single transition then endows state A with an RT of 280s. A2A’s and A2B’s state *i*, although they share a certain degree of similarity, present different interactions. While in A2A, NECA still presents the majority of the state A contacts, particularly for the ribose moiety and the 5’-N-ethylcarboxamide, this is not true for A2B. Therefore, state *i* in A2B is more similar to state *ii* of A2A. In A2A, the unbinding process seems to exhibit an extra intermediate state (state *i*) that A2B wholly skips. Essentially, A2B’s state *i* lost the optimized double H-bond contact with N^6.55^, it does not present an optimized *π*-*π* interaction with F173, and lost all the agonist-specific residues contacts with T^3.36^, S^7.42^, and H^7.43^; while it gained an interaction with the salt-bridge protagonists K267(H in A2A) and E174. This is a high-energy state where, as opposed to A2A, it does not present a high RT, and it transits to state *ii*, where it places itself at the interface between the OBS region and the vestibular site, completely losing all the native contacts with the OBS and interacting with the salt bridge. After which, NECA quickly exits to the unbound, solvent states. This low probability of transition between state *i* and *ii* is a phenomenon that can be appreciated from the FES presented in Figure 5, panel **b**. We notice that states *i* and *ii* are separated by a column of high-energy states in the range of CV distance between 1 and 1.4 nm. We rationalized this by noticing that there is an intricate interplay between the charged residues located in the extracellular lumen of A2B that is not present in A2A. In Fig. 5 **e** and **f**, we illustrate these differences. Essentially, panel **f** shows that there is an array of charged residues, namely D^1.32^(S in A2A), E174, K267(H in A2A), and K^7.32^(L in A2A), that function as a dynamic blockage for NECA. To visualize this, we generated a scatter plot that shows, on the x-axis, the distance between E174 and K267(H in A2A), on the y-axis, the distance between E174 and K^7.32^(L in A2A), and the scatter plot is colored in relation to the distance between NECA and TM1 from the structures corresponding to state *ii* in the FES. The plot is accompanied by a top view of the protein showing NECA in green and the aforementioned charged residues. What we observe is that there are three main types of unbinding conditions: (1) blue ones in which the distance between E174 and K267(H in A2A) is low, the distance between E174 and K^7.32^(L in A2A) is high, as the latter is in a salt bridge interaction with D^1.32^(S in A2A); (2) the green one in which none of the salt bridges between the protein are formed but there is a lipid insertion that does interact with D^1.32^(S in A2A); and (3) the red ones at which the distance between E174 and K267(H in A2A) is high and the distance between E174 and K^7.32^(L in A2A) is low forming a salt bridge. These conditions then have a consequential role in the unbinding pathway of NECA from state *ii*. From the distribution of points presented on the scatter plot in Fig. 5, **f**, we notice that the most sampled pathway is the red condition, in which NECA exits from the left side, where the steric hindrance of E174 is not present because it is interacting with K^7.32^(L in A2A), and thus it is the structure reported state in Fig. 5, **c**. Due to the higher density of charged residues on A2B’s extracellular side, their dynamics seem to have a non-negligible role in the egress of bulkier ligands like NECA. Something that we did not see on TEP due to its small size, nor on ZMA due to its phenol group that sits in the vicinity of K^7.32^(L in A2A), thus limiting its dynamics. There is something interesting to note about the blue and green states. If we look at the structural images below the scatter plot in Fig. 5, **f**, we see that NECA’s relative position to the protein is very similar between them; however, they are mapped to different TM1 distances on the scatter plot (in the blue ones, NECA is closer to TM1 than in the green ones). This difference in distance is not due to NECA’s absolute position, but instead it is due to an aperture of TM1 from the core TM bundle of the receptor. This change in the protein’s conformation is due to an insertion of a POPC lipid between TM1 and TM7. Interestingly, the positive choline group of the POPC directly interacts with D^1.32^(S in A2A) and the negative phosphate group could interact with K^7.32^(L in A2A). This lipid-protein salt bridge interlock results in the disruption of the salt bridge between D^1.32^(S in A2A) and K^7.32^(L in A2A), as seen in the blue states, and opens TM1 away from the protein core. Remarkably, the TM1-TM7 region has been noted to be a hotspot for lipid insertions in many Class A GPCRs, including A2A, in *in vitro* and *in silico* studies, suggesting that the insertion we observe here can have a physiological role.^106,107^ We observed this POPC insertion in In contrast, A2A does not seem to exhibit such a complex charge interplay. In Fig. 5, **e**, we plot a similar data set as we did for A2B, with the difference being that in A2A K267 is mutated to protonated H264, K^7.32^ is mutated to a non-polar L^7.32^, and D^1.32^ is mutated to an S^1.32^. Clearly, in A2A there is no dynamical interplay of charges in this region, and this leads to state *ii* exhibiting only one unbinding pathway with notably less charge and steric hindrance than in A2B. Thus, conclusively, this charged A2B extracellular region is responsible for the zone of high energy states in the FES shown in Fig. 5, **a** that separates the bound from the unbound states, and it is responsible for the high RT of NECA with its associated low probability of unbinding; two characteristics that are not present in NECA bound to A2A.

## Conclusion

In this study, we used an enhanced-sampling MD protocol to characterize the thermodynamics and kinetics of three A2A and A2B ligands: TEP, ZMA, and NECA. Thermodynamics were investigated using FM, which biases two intuitive geometrical CVs to efficiently sample the ligand–receptor phase space and reconstruct the corresponding FES. This approach reproduces the experimentally known binding modes of TEP, ZMA, and NECA at A2A and of NECA at A2B, while predicting the unresolved binding modes of TEP and ZMA at A2B. The calculated binding affinities also agree with experiment. Kinetics were characterized using AdInf, which biases the ligand–protein distance to generate multiple unbinding events and estimate residence times. Unbinding pathways were clustered into metastable states and represented through a transition matrix containing state-to-state residence times and transition probabilities. This kinetic model allowed us to identify the rate-determining step and the corresponding high-energy transition-state region. Together, FM and AdInf provide a complementary atomistic description of ligand binding thermodynamics and unbinding kinetics. Functionally, we suggest that the four point mutations found in the OBS site: L/V^6.51^, H264/K267, L/K^7.32^, and Y/N^7.36^ of A2A/A2B can have a defining role in the recognition, stability, and dynamics of these ligands in each of these receptors. For example, we found that the OBS on A2B exhibits a secondary pocket in the TM1, TM2, and TM7 region that is kept at a higher level of hydration when comparing the same region in A2A. We suggest that this difference is in part driven by the Y/N^7.36^ mutation. A higher number of water molecules in this region has a kinetic effect on the egress of ligands from these receptors, as was the case we observed with TEP. Moreover, the residue Y^7.36^(N in A2B) has been identified as necessary for the recognition of various ligands such as inverse agonist ZMA, the antagonist XAC, and the agonist CGS21680.^89,90^ Furthermore, the point mutation N^7.36^ in A2B has been identified to contribute 6-fold to the binding of the A2B-selective agonist BAY60-6583.^24^ Another example is the mutation L/V^6.51^, which has been experimentally found to be responsible for the recognition and affinity of ligands in A2B.^24^ We found that this mutation has a non-negligible effect on the volume of the OBS floor, especially when considering ligands that reach those deeper parts of the OBS, like ZMA. We suggest that the tighter VdW packing created by the OBS floor hydrophobic pocket formed of V^3.32^, L^3.33^, M^5.38^, H^6.52^, and L/V^6.51^ in A2A greatly stabilizes ZMA in the A2A OBS, effectively increasing its affinity, and is in part responsible for the higher RT observed in A2A. These observations suggest that A2A selectivity may extend beyond the exact triazolotriazine-furan scaffold of ZMA to antagonists sharing a similar fused bicyclic nitrogen-containing heteroaromatic core linked to a furan ring and adopting a comparable binding mode. Consistent with this hypothesis, compounds such as SCH442416, preladenant, and SCH58261 display higher reported affinity for A2A than for A2B despite variations in the identity of the bicyclic core, which all exhibit A2A selectivity. Finally, we also noted that the extracellular region of A2B contains a higher degree of charged residues when compared to A2A. This creates a dynamic interplay between the various charged residues, such as K, E, and D residues that are not present in A2A. Particularly, we found that the residues D^1.32^(S in A2A), E174, K267(H in A2A), and K^7.32^(L in A2A), all located at the extracellular lumen of A2B, can interchange their opposite-charge partners and have a significant role in the unbinding pathways and, therefore, the kinetic rates that bulkier ligands like NECA exhibit in this receptor. The A2A equivalent residues of this region are S^1.32^(D in A2B), E169, H264(K in A2B), and L^7.32^(K in A2B), which contain less net charge and less bulky residues when compared to A2B, and did not appear to have a direct effect on the pathway of NECA. The charged residue region correlation with the unbinding pathway was not observed in TEP-A2B or ZMA-A2B, possibly due to the small size of TEP and because the long functional phenol group of ZMA blocks these residues position into place when it is in its most stable binding pose. Moreover, this charged residue region, particularly between TM1 and TM7, was confirmed to be a functional lipid insertion, as has been confirmed by other studies.^106,107^ We conclude then that A2A and A2B, regardless of being two GPCR paralogs with a high degree of sequence identity, exhibit minute physical and chemical characteristics that endow them with the necessary differences to respond to different biological stimuli and environments in our bodies. These differences are difficult to grasp from static resolved structures and from *in vitro* studies. Here we suggest three OBS key characteristics that serve as a functional map when it comes to drug/lead design or drug repurposing for these two receptors: the solvation/polarity differences in the extracellular region of TM1, TM2, and TM7, volume differences in the at the bottom part of the OBS between TM3, TM5, and TM6, and charged residues presence and dynamics in TM1, ECL2, ECL3, and TM7 that dictate ligand pathways. More broadly, our findings establish a dynamic molecular foundation for understanding how subtle sequence variations can be amplified into distinct pharmacological profiles in closely related GPCRs. Incorporating such thermodynamic and kinetic determinants into ligand design may therefore provide additional opportunities to achieve subtype selectivity beyond those accessible from equilibrium structures alone, with direct implications for the rational development of more selective adenosine-receptor modulators.

## Acknowledgement

The results hereby reported are part of a project that has received funding from the European Research Council (Grant agreement No. 101001784 “CoMMBi”). This work was supported by a grant from the Swiss National Supercomputing Centre (CSCS) under project ID s1293 on Alps.

