## Supplementary Information for "Thermodynamic, Kinetic, and Structural Determinants of Ligand Selectivity in A2A and A2B Adenosine Receptors"

### Assessing GPCR Adenosine Receptors A2A and A2B Thermodynamics and Kinetics Using Advanced Computational Techniques

### Contents

**Sequence Alignment** **3**

**Modelling of the Extracellular Loop 2 in A2B** **3**

**CV Explanation** **6**

**Kinetics** **9**

### Sequence Alignment

With reference to Figure 1 in the main text, here we show the full sequence alignment of the two receptors.

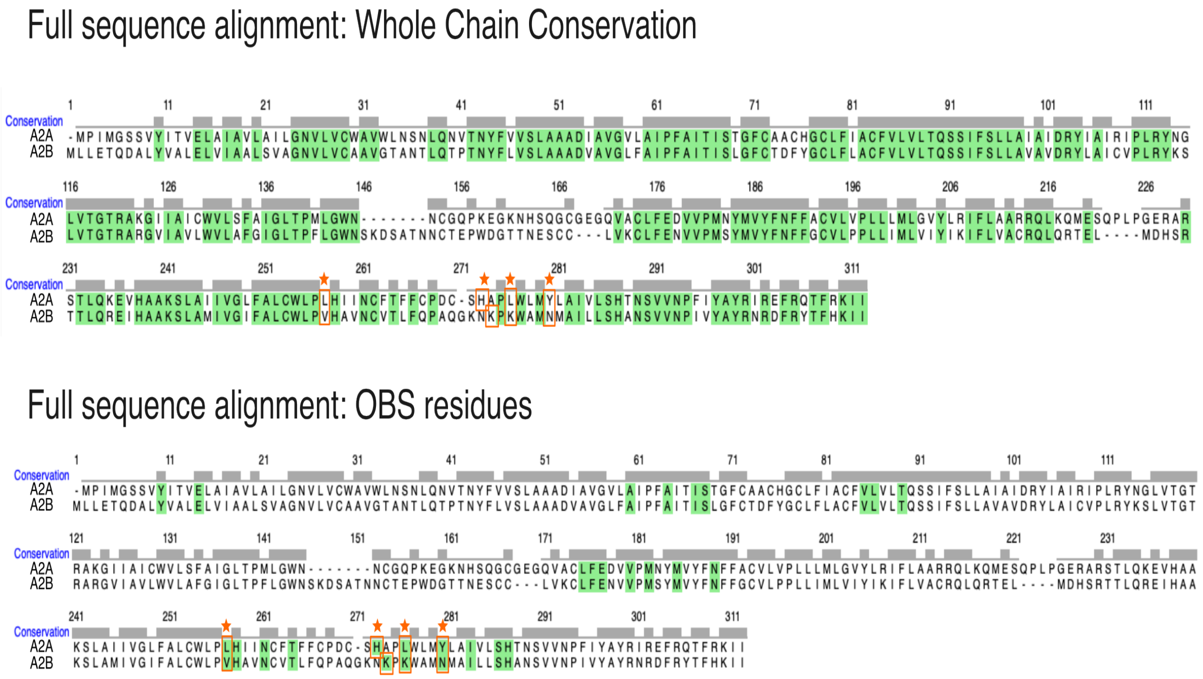

Figure 1: Sequence alignment done with the Clustal Omega tool in ChimeraX. In the top image, the highlighted green residues are conserved residues between A2A and A2B across the whole chain. We use this information to generate the color map of the structures shown in Figure 1 in the main text. Note that we have added four salmon colored stars to indicate the four point mutations that most concern the arguments presented in this study. On the bottom image, we then highlight in green the residues that form the OBS; most are conserved with the exception of the four indicated point mutations.

### Modelling of the Extracellular Loop 2 in A2B

The A2B's extracellular loop 2 (ECL2) was simulated while maintaining the cysteines present in this region (i.e., C154, C166, and C167) in their reduced state, as supported by experimental studies reported in the literature. Müller *et al.* show that in the A2B receptor, only the C78-C171 disulfide bond is essential for ligand binding and receptor activation.[?] Thus, the only disulfide bridge included in our structure is between C78 and C171. Given the limitations of state-of-the-art loop modeling algorithms in accurately predicting very long loops (longer than 20 residues, as in our case), we adopted a physics-based approach, combining MD simulations with enhanced sampling techniques.[?] In particular, we employed Replica-Exchange with Solute Tempering 2 (REST2), a Hamiltonian Replica-Exchange MD simulation method, using GROMACS 2021.5 patched with PLUMED 2.7.3, in order to simultaneously explore the conformational space of A2B's ECL2 and obtain an accurate starting structure for our thermodynamic and kinetics computations[?][?][?] The interaction potentials of intra-protein and protein-solvent were scaled by factors $\lambda$ and $\sqrt{\lambda}$, respectively, while water–water interactions were unaltered.[?] The scaling factor $\lambda_i$ , and corresponding effective temperatures $T_i$ of the $i^{th}$ replica are given by:

$$\lambda_i = \frac{T_0}{T_i} = \exp\left(-\frac{i}{n-1}\ln\left(\frac{T_{\max}}{T_0}\right)\right)$$

where $T_0$ and $T_{max}$ are the effective temperatures of the lowest-rank (unscaled) and the highest-rank replicas, respectively, and $n$ is the total number of replicas used. The "hot" region in our simulation encompassed the ECL2 region (residues S146 to V169, according to UniProt ID: P29275 numbering). A total of 12 replicas were performed, where the lowest temperature ($T_0$) was set to 300 K, and the highest temperature ($T_{max}$) was set to 600 K. Exchange of coordinates between neighboring replicas was attempted every 2 ps to reach an average exchange probability of $\sim$ 30%. Each replica of HREMD is $\sim$ 330 ns long, for an effective sampling time of $\sim$ 4.0 $\mu$s. For analysis, we use only the trajectory of the unscaled lowest-rank replica ($\lambda_0 = 1$ or $T_0$). Production runs were carried out in the NPT ensemble with a 2 fs timestep. Unless otherwise stated, all force fields and simulation parameters for energy minimization, system equilibration, and production MD are identical to those utilized for unbiased MD simulations in this study. To identify biologically relevant ECL2 conformations sampled in our HREMD simulations, we clustered the ECL2's essential subspace defined by Principal Component Analysis (PCA) of the unscaled replica ($T_0$), which describes opening and closing motions of ECL2. The "*sklearn.decomposition.PCA*" module was used to perform a PCA of the $C_\alpha$ in ECL2, while the "*sklearn.cluster.DBSCAN*" module was used to implement the DBSCAN (density-based clustering for applications with noise) clustering algorithm in order to identify clusters as regions of high point density in the essential subspace. The centroid of the most populated cluster was used as a starting structure for subsequent A2B simulations.

#### CV Explanation

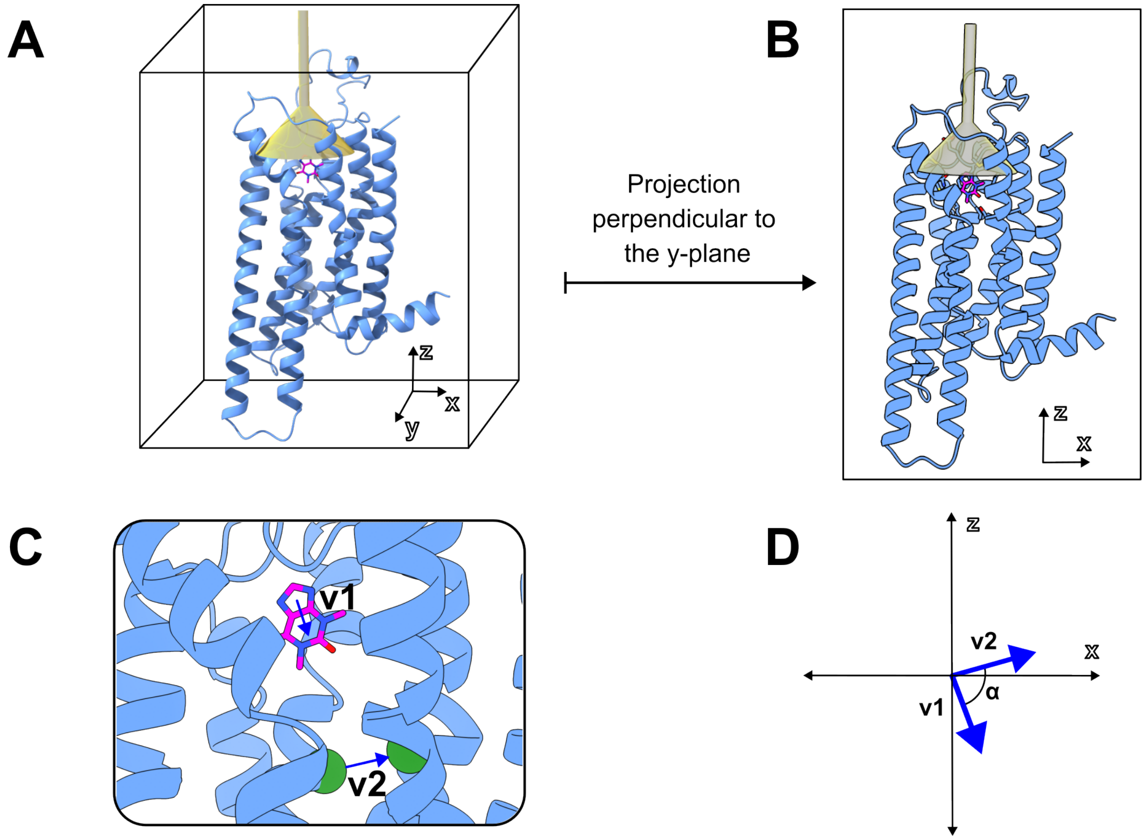

Figure 2: (A) Shows the three-dimensional structure of inactive A2A in complex with theophylline, along with a representative placement of the funnel used in the FM calculations. The protein is represented in blue ribbons, theophylline is represented in magenta sticks, and the funnel is colored transparent yellow. (B) Shows the projection of the torsional CV vectors from the three-dimensional space into the x,z-plane. (C) Shows a detailed representation of theophylline in A2A's OBS, along with the two vectors used for the torsional CV calculation represented as arrows. **v1** is a vector that is defined by the COMs of the two rings of the xanthine moiety of the adenosine receptor ligands. Vector **v2** is defined by the position of the $C_\alpha$ of specific residues located in helix 2 and helix 3. The selected $C_\alpha$ atoms are shown in green spheres. (D) Shows a schematic illustration of the angle $\alpha$ formed by the two projected vectors **v1** and **v2** that are accelerated throughout the FM calculation.

In panel A of the figure, a representation of the funnel placement is shown, which covers the entrance to the protein's OBS. We want to emphasize that the simulation is defined in the three-dimensional Euclidean space, where each point has an associated (x,y,z) coordinate. However, the torsional CV that we chose to define and accelerate throughout all of our FM calculations projects the ligand's rotational degree of freedom into a two-dimensional plane that is parallel to the planar xanthine rings of the ligand. This projection is conveyed in panel B of the same figure. On panel C, we then show in detail TEP bound to the A2A's OBS, in which we illustrate the two vectors that compose the torsion, namely, **v1** and **v2**. Vector **v1** is defined as a vector that connects the two centers of mass of the xanthine moiety rings, shared by all the ligands that we considered. Special care is taken for the ribose-containing ligands, such as NECA. In all cases, **v2** is a vector defined by two stable $C_\alpha$ atoms in the protein, which in our systems are located in the center of TM helix 2 and TM helix 3. Having a reference vector **v2** defined by the protein has the beneficial effect that it becomes an inertial frame of reference onto which we can track and measure the ligand's rotation, maximizing the rotational CVs' injectivity.

As discussed in the Methods section of the Main Text, we performed the FM simulations biasing two geometric CVs: (1) the distance between the center of mass of the ligand and the binding pocket of the protein and (2) the torsional degree of freedom projected in the x,z-plane of the simulation space (See Fig. 2 of the Main Text). The ligand vector (labeled **v1** in Fig. 2 of the Main Text) is defined in a similar way for TEP and ZMA ligands. Instead, as introduced in the Main Text, we had to slightly redefine this vector for NECA. The ribose moiety of NECA introduces larger degrees of freedom that would not be accounted for using the **v1** vector used for TEP and ZMA. Therefore, we decided to modify the torsional vector for NECA in such a way that it includes the rotational degree of freedom of the purine scaffold together with the ribose moiety. In Supplementary Fig. 3 panel a, we illustrate the vector as defined in our thermodynamic and kinetic simulations. Instead, in panel b, we show what would happen if we use the same definition of **v1** in TEP and ZMA in NECA. Note that the vector in panel b is not able to capture and differentiate completely different binding poses, since it completely ignores the ribose orientation.

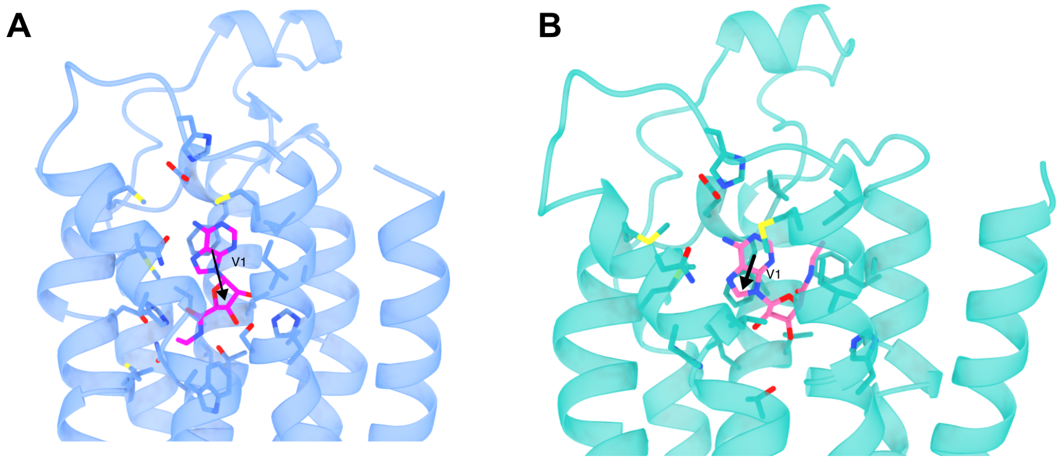

Figure 3: **a)** Torsional vector **v1** used for the NECA simulations. The two points that define the vector are the center of mass of the purine moiety and the center of mass of the ribose moiety. A2A and NECA are represented by blue ribbons and magenta sticks, respectively. **b)** Torsional vector **v1** as defined for TEP and ZMA, applied to NECA. As is evident from the figure, this torsional angle would be completely ignorant of the ribose moiety and thus would not differentiate the binding modes in the CV space. A2A and NECA are represented in green ribbons and magenta sticks, respectively.

### Kinetics

##### A. State-to-state rates and Transition Matrices

In this section, we present the heatmaps generated by mapping all the unbinding events into the distance-torsion CV space. As specified in the Main Text, we use these heatmaps to define the metastable states of the ligand while it unbinds and build the transition matrix $M$. Here we label the hot spots of interest with Arabic numbers. These numbers then correlate with the states in the transition matrix $M$. In Supplementary Figures 4 and 6, we present the heatmaps for all three ligands as they unbind from A2A and A2B, respectively. On the right side of each heatmap, we show the transition matrix using the metastable state definitions shown in the heatmaps.

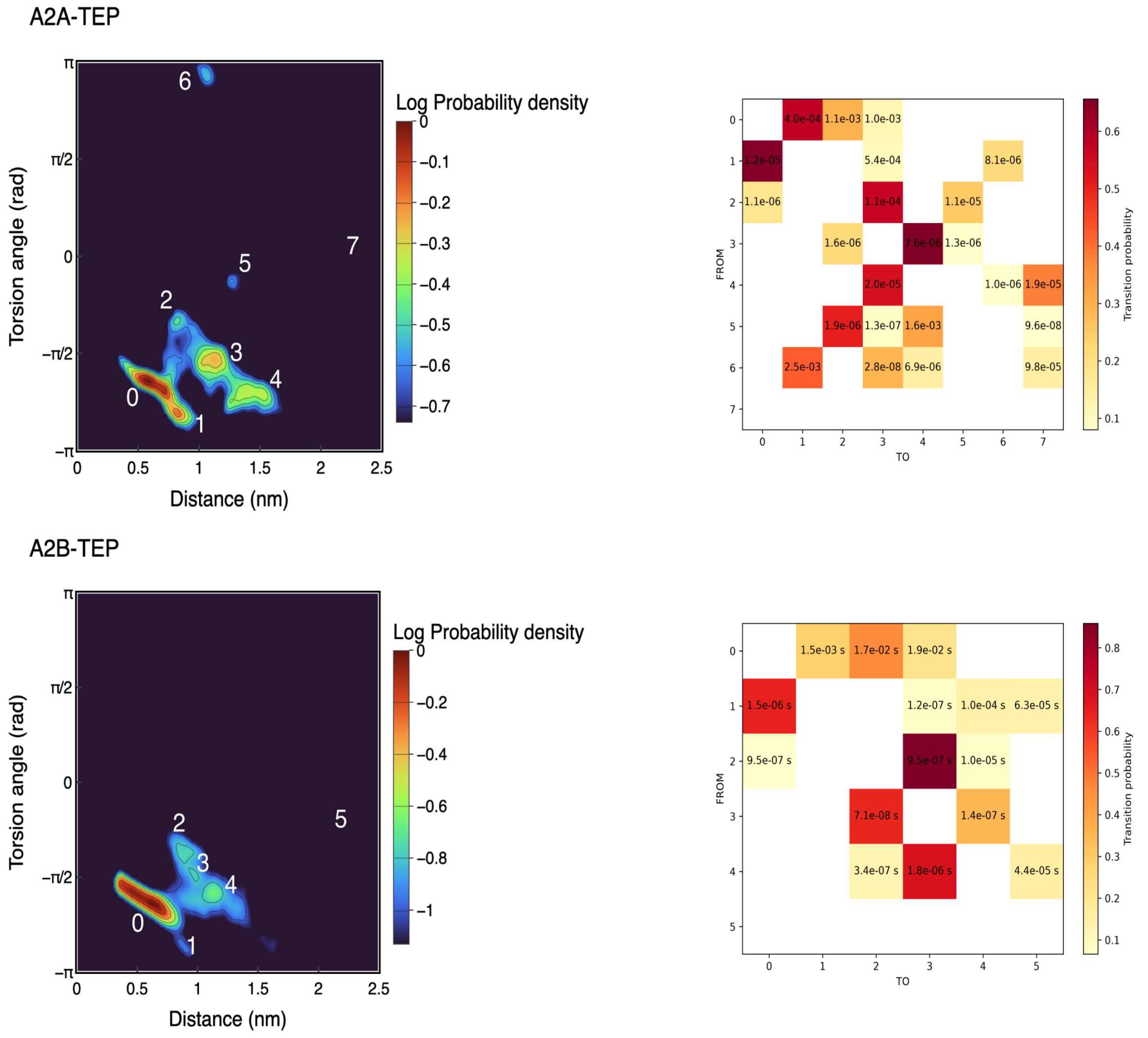

Figure 4: Heatmaps of TEP, ZMA, and NECA as they unbind from A2A, projected in the Torsion-Distance CV space. The heatmaps are normalized and colored by the number of counts in each bin. The numbers define the metastable states we identify in the heatmap plot. The vertical white line indicates the threshold at which no more features are shown in the heatmap; therefore, we consider the ligand unbound (the highest number in our definitions). On the right side of each heatmap, we show the full transition matrix $M$, which we use to analyze the transitions reported in the Main Text. Each element $M_{ij}$ in the matrix indicates the RT and the probability of belonging to the transition from $i$ to $j$. The color scale indicates the probability, and the number written on each matrix element is the RT in seconds.

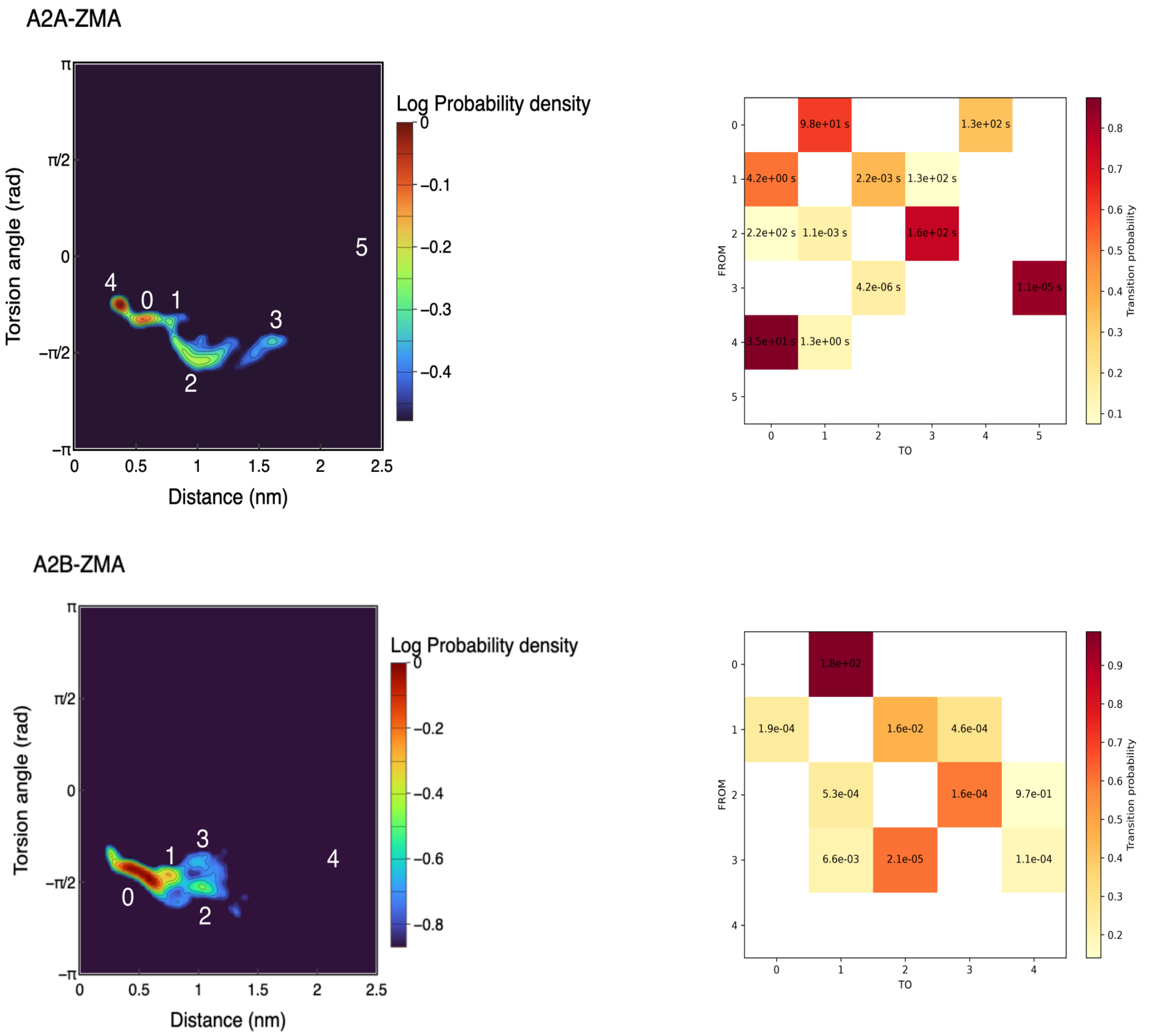

Figure 5: Heatmaps of TEP, ZMA, and NECA as they unbind from A2B projected in the Torsion-Distance CV space. The heatmaps are normalized and colored by the number of counts in each bin. The numbers define the metastable states we identify in the heatmap plot. The vertical white line indicates the threshold at which no more features are shown in the heatmap; therefore, we consider the ligand unbound (the highest number in our definitions). On the right side of each heatmap, we show the full transition matrix $M$, which we use to analyze the transitions reported in the Main Text. Each element $M_{ij}$ in the matrix indicates the RT and the probability of belonging to the transition from $i$ to $j$. The color scale indicates the probability, and the number written on each matrix element is the RT in seconds.

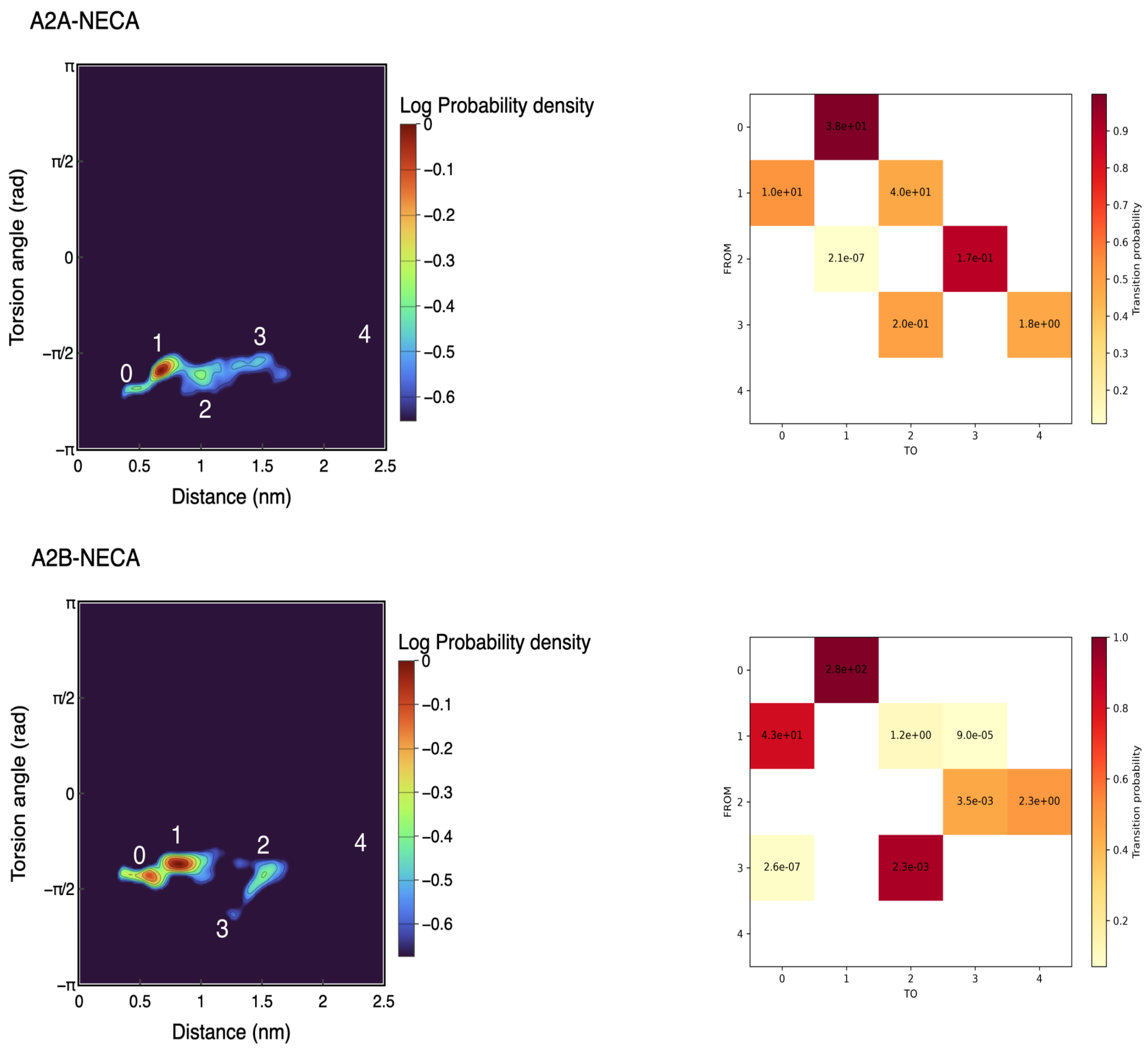

Figure 6: Heatmaps of TEP, ZMA, and NECA as they unbind from A2B projected in the Torsion-Distance CV space. The heatmaps are normalized and colored by the number of counts in each bin. The numbers define the metastable states we identify in the heatmap plot. The vertical white line indicates the threshold at which no more features are shown in the heatmap; therefore, we consider the ligand unbound (the highest number in our definitions). On the right side of each heatmap, we show the full transition matrix $M$, which we use to analyze the transitions reported in the Main Text. Each element $M_{ij}$ in the matrix indicates the RT and the probability of belonging to the transition from $i$ to $j$. The color scale indicates the probability, and the number written on each matrix element is the RT in seconds.

#### B. Kolmogorov-Smirnov tests

In this section, we present the Kolmogorov-Smirnov (KS) test applied to our unbinding times for each ligand-receptor pair. In Supplementary Figure 7, we show the fit of the theoretical exponential cumulative distribution function with our computed cumulative distribution function for all ligands.

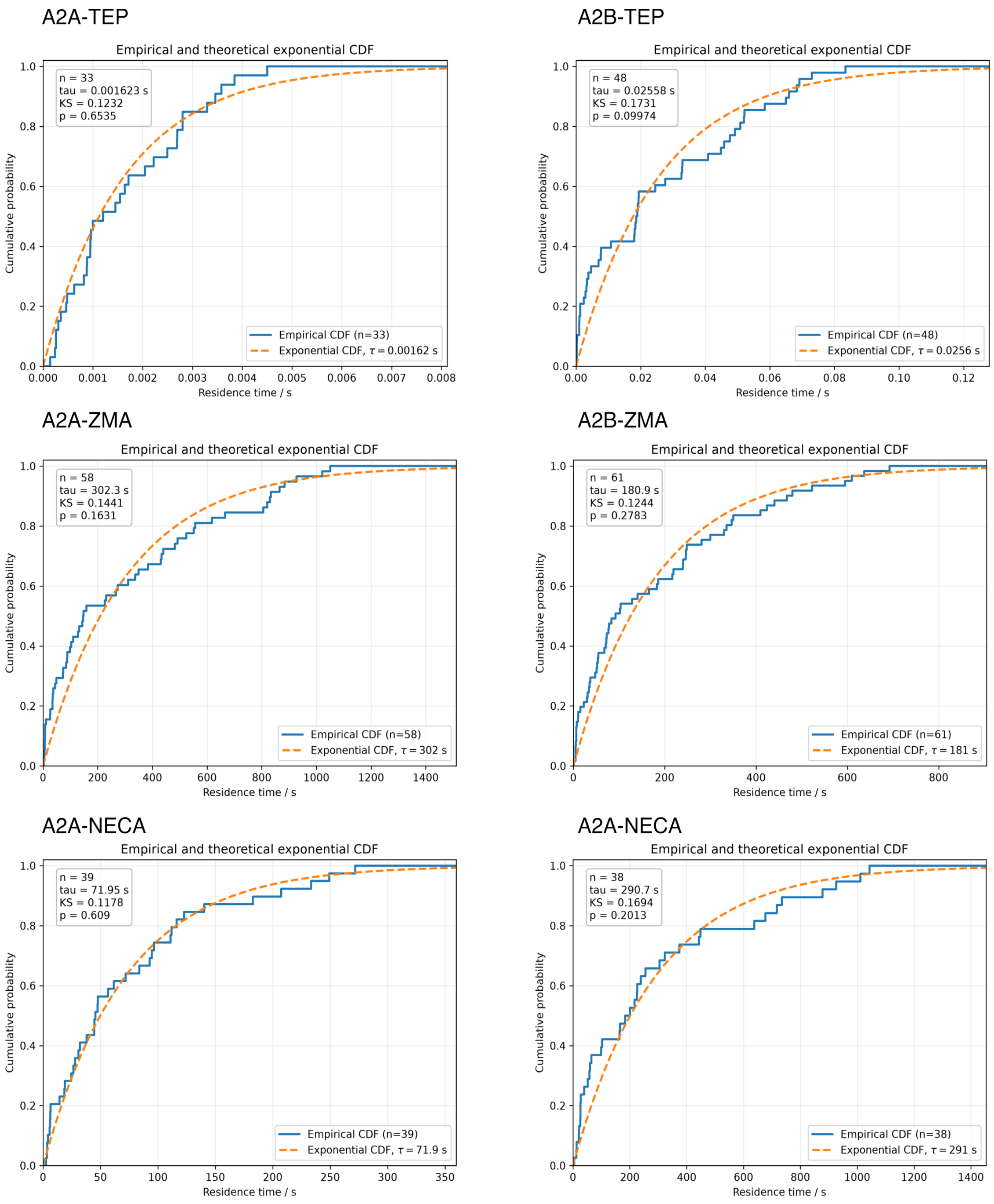

Figure 7: KS test for all ligand-protein complexes when tested against the theoretical (blue line) and empirical cumulative distribution functions (orange line). Below each title, we report the two metrics obtained from the KS test: the distance between the distributions, $D$, and the *p-value*.

### Unbiased MD Simulations of Thermodynamic Minima

#### A. Ligand-RMSD and CV Fluctuations

In the following section, we show the results of the unbiased MD simulations using as a starting structure a frame obtained from the thermodynamic minimum of each FM run. In Supplementary Fig.8 we show the CV (distance and torsion angle) and RMSD values for the three ligands in complex with A2A. In Supplementary Fig.9 we show the same analyses for the ligands in complex with A2B.

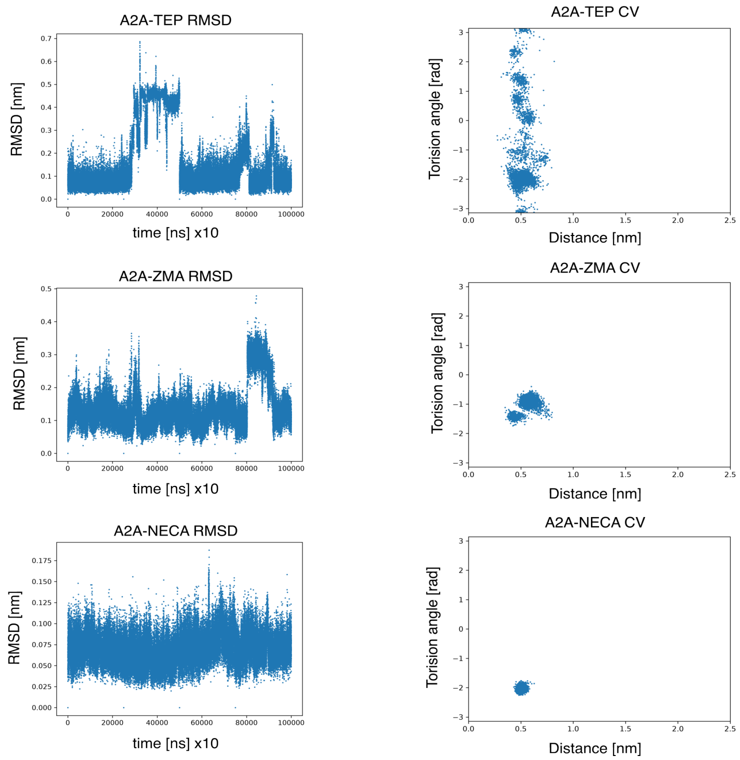

Figure 8: On the left column of the figure, we show the RMSD with respect to the initial frame of the simulation as a function of simulation time. On the right side of the figure, CV values of the ligands in the unbiased MD simulation in complex with A2A are reported.

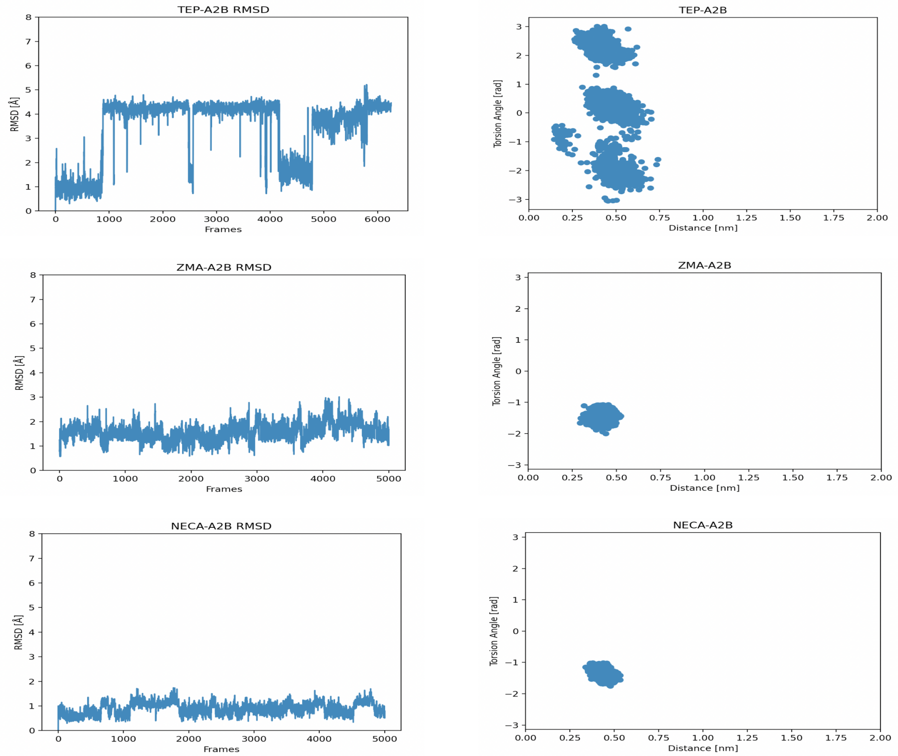

Figure 9: In the left column of the figure, we show the RMSD with respect to the initial frame of the simulation as a function of simulation time. On the right side of the figure, CV values of the ligands in the unbiased MD simulation in complex with A2B are reported.

#### B. Orthosteric Binding Site Volume

In this section, we show the available volume in the region corresponding to the hydrophobic pocket deep in the OBS of the A2A and A2B receptors (Supplementary Fig. 7). This pocket is formed by the residues: L85/86, V84/85, L249/V250, and H250/251 in A2A/A2B respectively. As mentioned in the main text, there is an appreciable difference in volume, which could be due to the L/V mutation between the receptors. As discussed in the Main Text, this difference in volume could have a stabilizing effect on the apolar moiety of ligands that reach these deeper parts of the pocket, such as ZMA. The volume is calculated by using an in-house script that measures volumes in a similar fashion to how POVME works. We defined an inclusion sphere whose center is defined by the center of mass of the residues V85/86 and L249/V250 (A2A/A2B), with a 4 Å radius. The output is a .dx file that is essentially a three-dimensional grid which we can visualize in ChimeraX by selecting different iso-surface values that range from zero to one. The iso-surfaces have a physical meaning. Iso-surface values that are close to zero mean that they are spatial points that are transiently available and not really available volume that a ligand (ZMA in this case) can use to its entropic favor. On the opposite end, iso-surfaces close to one mean that they are spatial points that are consistently available throughout the simulation, and thus they represent space not occupied by the protein at any time. Values between these two extremes then follow this logic in a proportional manner. In Supplementary Fig. 7, we show the A2A and A2B orthosteric binding pocket volume represented by the iso-surface with value 0.2 obtained from our calculations using an unbiased 1 $\mu$s simulation for the receptors coupled to ZMA. We found it informative to compute the volume of these iso-surfaces using ChimeraX's 'blob' module and plot them as a function of increasing iso-surface value. We make note of the iso-values that lie between 0.2 and 0.9, which represent the accessible volume of the receptor's orthosteric pocket. In all of this range of values, A2A's pocket volume is lower than that of A2B. Since the only variable that changes between these two measurements is the point mutation L/V, we then attribute this substantial difference to the longer side chain of L in A2A. Alternatively, we computed the volume variations throughout the simulation and generated a cavity volume density. This plot also confirms that A2A's volume is generally smaller than for A2B. This, as discussed in the main text, gives rise to all sorts of entropic and enthalpic effects on the ligand binding of ZMA, and we suppose, on the binding of similar ligands on these two receptors.

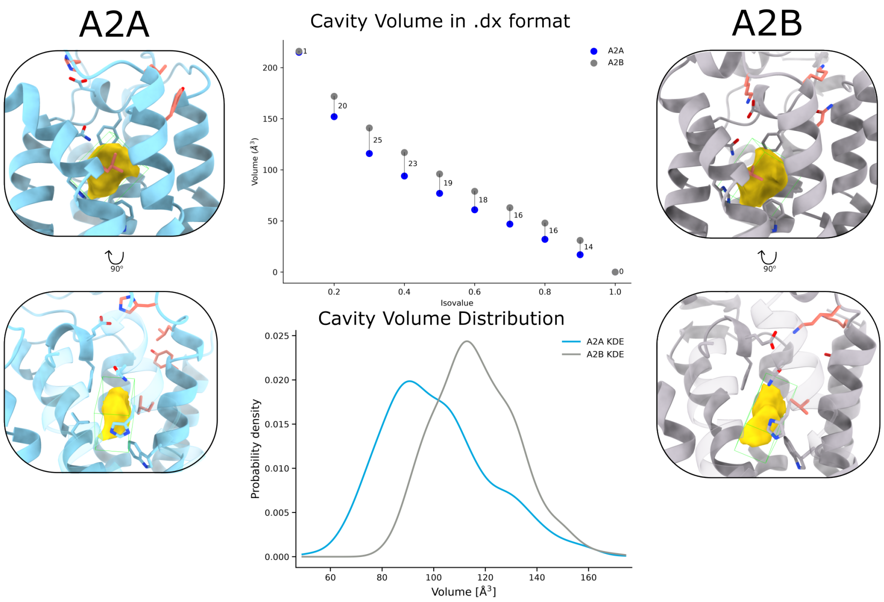

Figure 10: Volume of the hydrophobic pocket formed by the residues L85/86, V84/85, L249/V250, and H250/251 in A2A/A2B respectively. The average volume iso-surface with value 0.2 within the OBS of A2A and A2B receptors is shown as a gold surface. On extremes left and right sides of the figure, we show A2A and A2B, respectively. In the center top part of the figure, we show the available cavity volume as computed using ChimeraX's 'blob' module as a function of iso-surface values. Each scatter plot value is the 'blob's volume in each receptor, and the vertical line with values is the difference in volume at each isovalue level. In the center bottom of the figure, we show the distribution of cavity volume as sampled from the unbiased simulation.

#### C. $Y/N^{7.36}$ Hydration Gate

As discussed in the main text, we observe a difference in the OBS solvation due to the $Y/N^{7.36}$ mutation between the two receptors. Due to the difference in size and the different electronic properties between these two residues, A2B's OBS tends to have a higher density of water molecules than A2A. In Fig. 11, we show the distribution density of water molecules around 5 Åof the centroid defined by the $C_\alpha$ atoms of Y/N$^{7.36}$ and S$^{2.65}$. As the figures suggest, there is a larger solvation shell around N than around Y. This occupied volume of water molecules is computed using an in-house algorithm similar to VMD's VolMap tool. We compute the average 3D position of oxygen atoms within the inclusion sphere defined above, and create a .dx file with this information. We then upload the structure and the

.dx file to ChimeraX, where we show both .dx volumes at the same isovalue (0.015 in this case) and compute their volumes using the 'blob' module. This gives 144.78 Å$^3$ to A2A and 333.86 Å$^3$ to A2B. Furthermore, below the figures, we show a smoothed histogram that shows the number of water molecules that are within this same inclusion sphere using PLUMED's COORDINATION module. This particular histogram is computed using the unbiased simulation of TEP in each receptor. The histograms confirm that A2A's amount of water in this region is lower than in A2B, with means of 7 water molecules in A2A and 11 water molecules in A2B. The histogram has a 'spiky' look because the water count is discrete, and we perform kernel density smoothing to ease visualization.

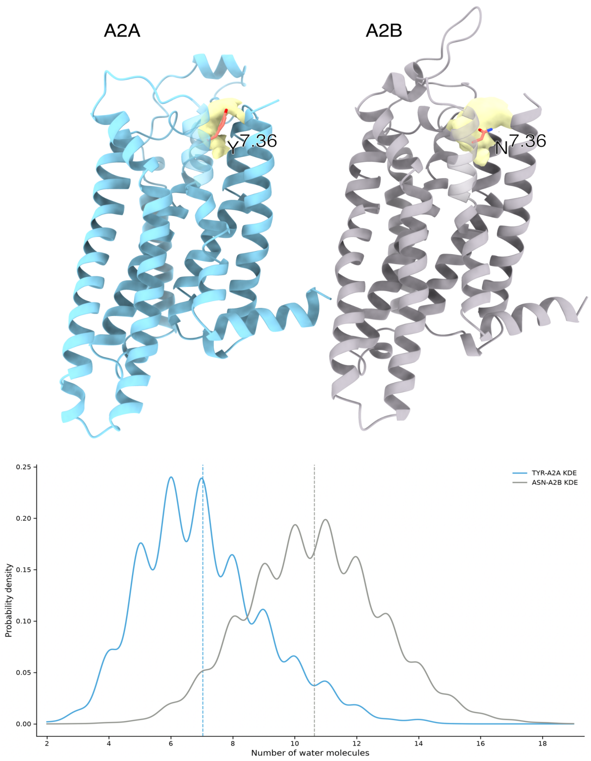

Figure 11: Water molecule presence comparison in the hydration gate pocket formed by $Y/N^{7.36}$. On the top part of the figure, we show the average spatial representation of water molecule presence in an inclusion sphere defined by the centroid of the $C_\alpha$ atoms of $Y/N^{7.36}$ and $S^{2.65}$. The volumetric information for both receptors is shown as a yellow surface at the iso-surface value of 0.015. The shown volume occupies 144.78 $\mathring{A}^3$ in A2A and 333.86 $\mathring{A}^3$ in A2B. In the bottom part of the figure, we compute the number of water molecules in this inclusion sphere for an unbiased simulation of TEP in both receptors and plot the distribution. Both of these measurements, volume and water count distributions, confirm the fact that A2A contains fewer water molecules on average than A2B within this hydration gate region.
